# Atomic-scale mechanisms of GDP extraction by SOS1 in KRAS-G12 and KRAS-D12 oncogenes

**DOI:** 10.1101/2024.06.17.599303

**Authors:** Zheyao Hu, Jordi Martí

## Abstract

The guanine exchange factor SOS1 is a crucial node into the positive feedback regulation of the KRAS signaling pathway. Currently, the regulation of KRAS-SOS1 interactions and KRAS downstream effector proteins has become a new hotspot in the development of KRAS-driven cancer therapies. However, the detailed dynamic mechanisms of SOS1-catalyzed GDP extraction and the impact of KRAS mutations remain unknown. Herein, the main mechanisms of GDP extraction from KRAS oncogenes by means of the guanine exchange factor SOS1 are disclosed and described with full details at the atomic-level. For GDP-bound wild-type KRAS, four amino acids (Lys811, Glu812, Lys939 and Glu942) responsible for the catalytic function of SOS1 were identified. With the occurrence of KRAS-G12D mutation, the GDP extraction rate is significantly increased. The molecular interactions behind this phenomenon have been subsequently identified being mainly hydrogen bonding interactions between the mutated residue Asp12 and a positively charged pocket located at the intrinsically disordered region^807−818^ and composed by Ser807, Trp809, Thr810 and Lys811. Our findings provide new insights into the SOS1-KRAS interactions and facilitate the development of related anti-cancer strategies based on the blockage of the above described mechanisms.

## 1. Introduction

The study of the RAS family of oncogenes is a hot topic, mainly due to their active participation in about one third of human tumours^1, 2^. RAS functions as the molecular switch, transmitting biological signals by cycling between inactive (GDP-bound) and active (GTP-bound) states^3–8^. RAS coordinates intracellular signaling cascades and regulates cell survival and proliferation by interacting with effector proteins. The dysregulated RAS signaling ultimately leads to cancer^9–11^. Kirsten RAt Sarcoma (KRAS) is one of most frequently mutated oncogene in several human tumors, such as lung^12^, colorectal^13^ and pancreatic^14, 15^ cancers. Among them 66% of KRAS mutations occur at its codon 12^16^. In the last decades intense research has been devoted to find methods able to block the growing of KRAS and its-mediated tumor^17–22^. Only two drugs have been approved by the U.S. Food and Drug Administration up today, both of them related to the KRAS-G12C mutation^23^: AMG510 sotorasib^24, 25^ and MRTX849 adagrasib^26^. In the case of the elusive but more prevalent and common KRAS-G12D mutation^27^, two compounds (MRTX1133^20^ and TH-Z835^28^) have been proposed but none has been authorized yet.

Setting aside the high difficulty of directly targeting strategies, indirect ways have shown great potential^29^. Several attempts have been done to indirectly target the KRAS-driven cancers through KRAS associated downstream effectors, such as targeting the members of RAF-MEK-ERK cascade^30–32^ and PI3K-AKT-mTOR signaling pathway^32, 33^. Moreover, the regulation of KRAS GDP-GTP cycle also shows its unique anti-cancer potential. The balance of active and inactive states dynamically controls the extent and kinetics of the KRAS associated signaling^34–37^. Targeting Son of Sevenless (SOS1)-KRAS interactions and KRAS effectors alone or in combination has shown promising prospects of against KRAS-driven cancers^38–42^. Therefore, the detailed mechanisms of SOS1 activation of GDP-bound KRAS species is of crucial importance for the development of new cancer treatment strategies. SOS1 is one of the most important guanine exchange factors (GEF) for the positively regulation of KRAS GDP-GTP balance, which catalyzes the GDP-bound KRAS at its catalytic binding site (CDC25, residues 780–1019) and promotes the dissociation of GDP^43, 44^. In addition to the CDC25 site, the RAS exchanger motif (REM, residues 567-741) of SOS1 can enhance the binding of GTP-bound KRAS thereby promoting its guanine exchange function^45–49^. The interaction interface of nucleotide-free HRAS bound to the CDC25 has been revealed by co-crystallization of HRAS and SOS1^50^. However, the atomic-level molecular mechanisms of CDC25-catalyzed GDP extraction and the impact of KRAS mutations on such process remain unknown.

In this work, we have employed microsecond-scale molecular dynamics (MD) simulations to model and monitor two species of GDP-bound KRAS oncogenes (wild-type and G12D) under interaction with SOS1. For each GDP-bound KRAS species, we performed three independent simulations and investigated the molecular interaction mechanisms and their corresponding time-scales at the atomic-level. In the GDP-bound wild-type KRAS case, the intrinsically disordered regions (IDRs^807−818^, residues 807-818) together with *α*-helix H (renamed as *α*H, residues 930-944) participate in the GDP extraction process. In particular, SOS1’s residues Lys811, Glu812, Lys939 and Glu942 play a major role in the catalyzed KRAS-wild type GDP release. With the occurrence of KRAS-G12D mutation, the synergy of IDRs^807−818^ and *α*H is well-conserved, while the negatively charged Asp12 can be captured by the positively charged pocket composed of Ser807, Trp809, Thr810 and Lys811, creating a more compact KRAS-SOS1 interaction interface. These small changes result in significant enhancement of the GDP extraction rate. Finally, the findings reported in this work provide a clear description of GDP extraction mechanisms with atomic precision, so that the positively charged pocket located on the IDRs^807−818^ region can be considered as a potential target to interrupt the interactions between mutated KRAS and SOS1, leading to effective blocking of GDP extraction and the consequent reduction of KRAS-driven cancers.

## 2. Results

### 2.1 Structure of KRAS-GDP-SOS1 complexes

With the aim to investigate the role of SOS1 as a guanine exchange factor and, especifically, to decipher how the CDC25 catalytic site of SOS1 promotes the guanine exchange of KRAS, we performed all-atom microsecond scale MD simulations of the KRAS-SOS1 complex in solution. To abbreviate the system notations throughout the text, we introduce two superscripted letters on KRAS. In particular, KRAS^*WT*^ and KRAS^*G*12*D*^ denote wild-type KRAS and the KRAS-G12D mutation, respectively. The simulations were performed for KRAS-SOS1 dimers, with KRAS interacting with SOS1 at the CDC25 catalytic sites. A total amount of 6 independent trajectories of lenght 4.2-6 *µ*s each were simulated and analyzed. In our simulations, SOS1 is composed of REM and CDC25 parts (see Fig.1A). As for KRAS, since the C-terminal hypervariable region (HVR) of KRAS is disordered and not involved in the RAS-SOS interactions, the highly conservative G-domain of KRAS is employed (see Fig.1B). Previous investigations^43, 44^ suggested that the CDC25 catalytic domain is responsible for the catalytic activity of SOS1, while REM plays an auxiliary role: the activated RAS at the allosteric site helps accelerate the rate of GDP extraction happened on the CDC25 catalytic site^45–49^. To better understand the role that CDC25 plays in guanine exchange and to exclude the acceleration effect of REM on CDC25 functions when comparing the impact of KRAS-G12D mutation on nucleotide exchange, we adopted the KRAS-SOS1 dimer as our main research subject, in which the GDP-binding KRAS was superimposed onto the CDC25 catalytic site of SOS1 via protein-protein docking powered by ClusPro server^51–54^ (see Fig.1C).

**Figure 1.**
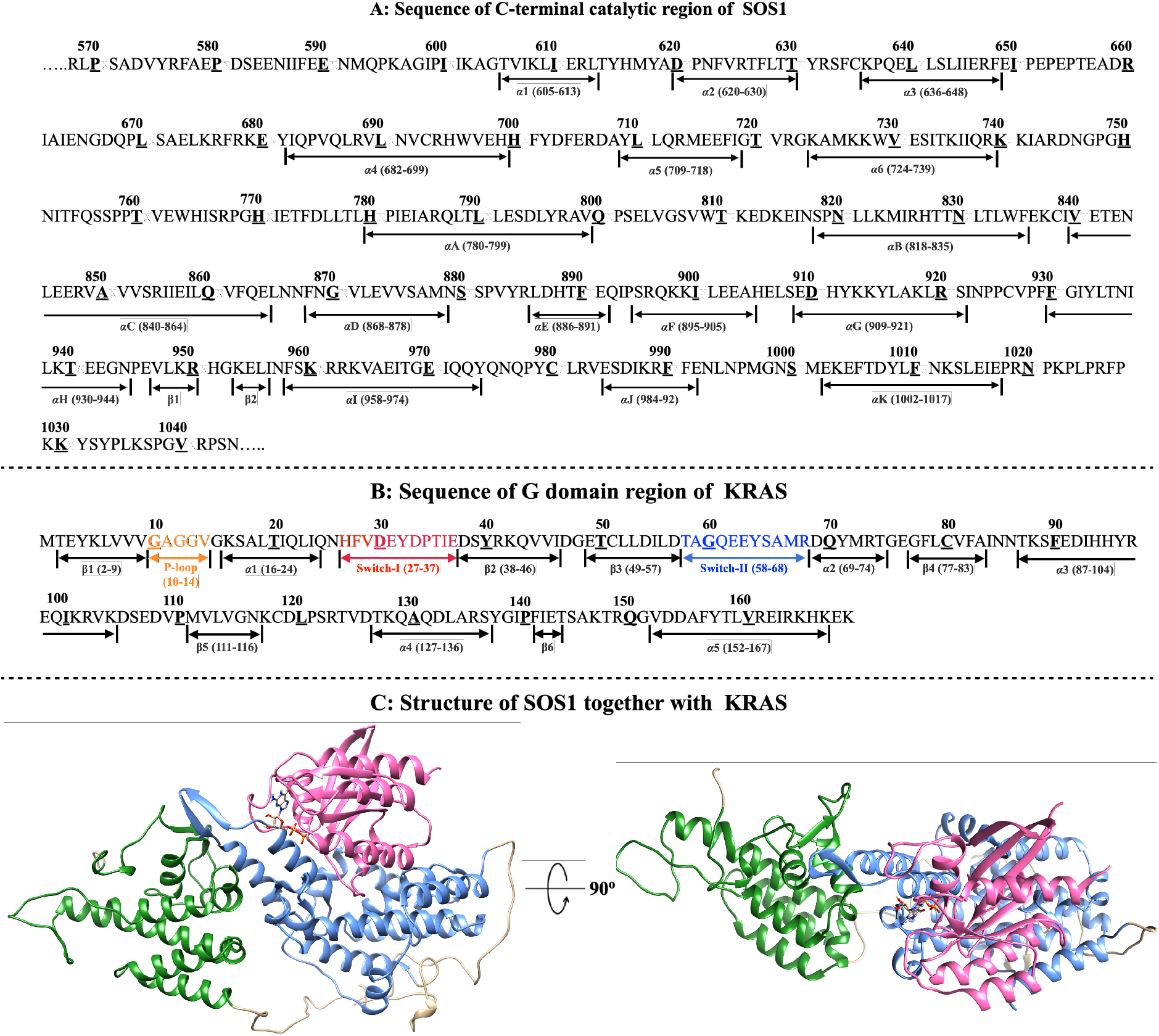
The structure of KRAS-GDP-SOS1 complexes. (A) The amino acid sequence of the C-terminal catalytic region of SOS1 formed by REM (residues 567-741) and CDC25 (residues 780–1019) parts; (B) The amino acid sequence of the catalytic domain of KRAS; (C) The complex SOS1-GDP-KRAS. REM and CDC25 domains are colored “forest green” and “cornflower blue”, respectively, whereas KRAS is colored “hot pink”. GDP is shown in “licorice” representation.

### 2.2 Molecular interactions of GDP extraction: wild-type KRAS case

As the first case, we studied the wild-type KRAS gene interacting with SOS1. We have observed that the molecular interactions by which SOS1 promoted KRAS nucleotide exchange can be mainly divided into the three consecutive stages: (1) preparatory stage (State-A, see Fig.2A), (2) molecular interaction induction stage (State-B, see Fig.2B) and (3) post-GDP extraction stage. Firstly, the preparatory stage mainly carries out the preparation process for GDP extraction, which is mainly reflected in the loading of KRAS^*WT*^ into the CDC25 catalytic site and the fine-tuning of the KRAS^*WT*^ -SOS1 interaction interface. The time scale of GDP extraction “State-A” is 1.5 ± 0.3 *µ*s. During this stage, the IDRs^807−818^ and *α*H regions gradually approach the KRAS^*WT*^ -GDP complex, as indicated in Fig.2 G-”State-A”. Among them, IDRs^807−818^ mainly point to the P-loop/GDP region of KRAS^*WT*^ and the main function of *α*H is to cleave and insert itself into the switch-I (SW-I) domain of KRAS^*WT*^ (see correlation between SW-I and the KRAS^*WT*^ main body at Fig.2 D). Secondly, the molecular interaction induction stage holds the tightest KRAS^*WT*^ -SOS1 interaction interface and, more likely, the “rate-limiting step” of SOS1-catalyzed KRAS nucleotide exchange process, where SOS1 competes with KRAS^*WT*^ for GDP binding priority. This conformational evolution requires a longer time, at the scale of 2.9 ± 0.1 *µ*s (see Fig.2 G-”State-B”.) During “State-B”, the IDRs^807−818^ and *α*H region of SOS1 are mainly responsible for GDP extraction mechanisms. Among them, residues Lys811, Glu812, Lys939 and Glu942^55^ of SOS1 are the most important functional amino acid residues. The positively charged residues Lys811 and Lys939 form a stable salt bridge interaction with the negatively charged phosphate group of GDP through electrostatic interactions. Polar amino acids Glu812 and Glu942 form stable interaction with the guanosine and pentose sugar ribose structure of GDP through hydrogen bonding interactions respectively (see Fig.2 B and E). Finally, GDP is detached from the guanosine binding site of KRAS^*WT*^ through the conformational evolution of “State-B”. During the post-GDP extraction stage GDP can be captured by the temporary GDP-binding pocket on the SOS1, which is mainly composed of IDRs^807−818^ and the *α*H domain (as indicated in Fig.2 C and F).

**Figure 2.**
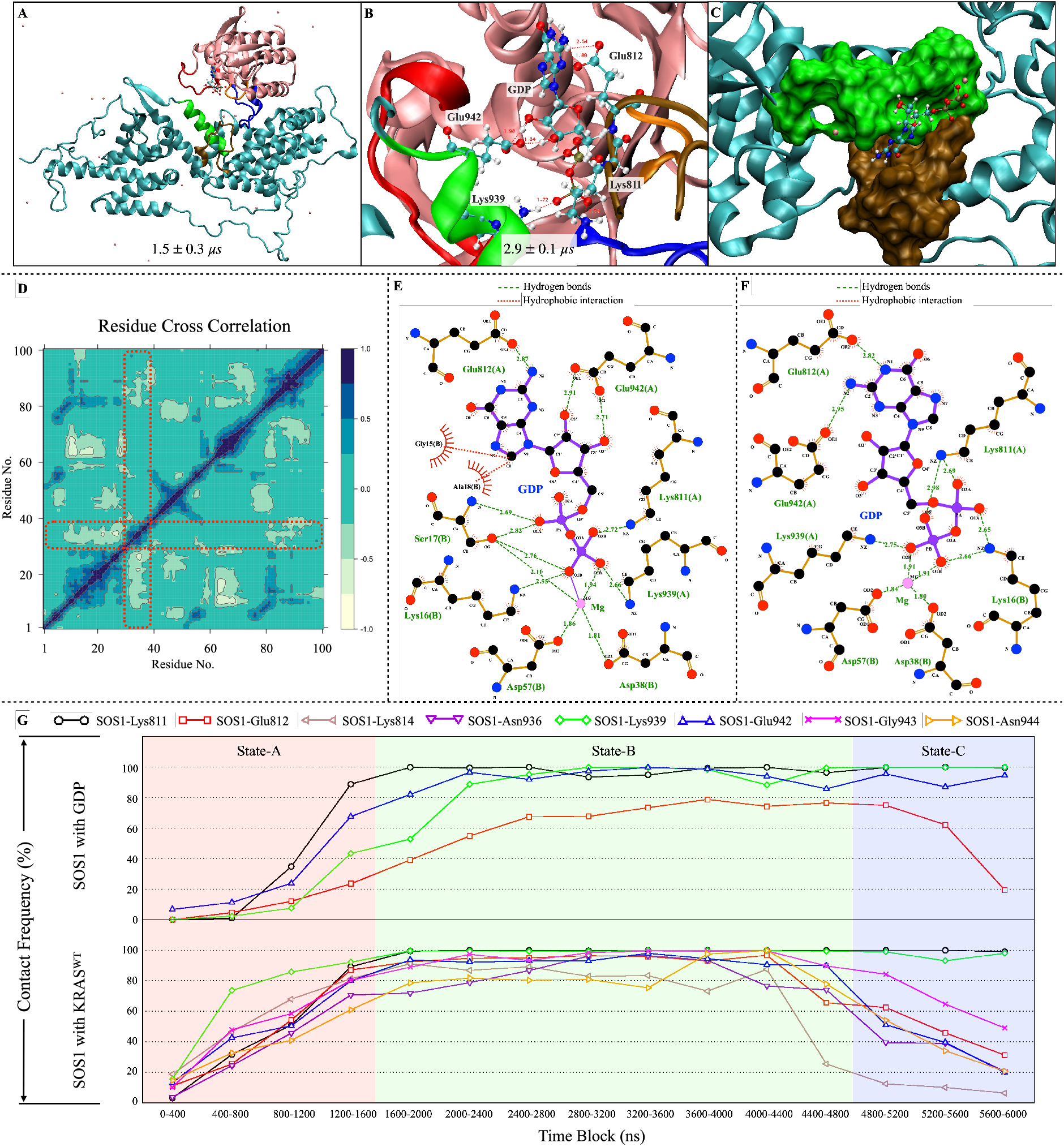
The molecular interactions of GDP extraction process for the KRAS^*WT*^ -GDP-SOS1 trimer. (A) State-A: Preparatory stage of GDP extraction. The schematic diagram of the KRAS^*WT*^ -SOS1 interactions in the preparation stage, and the time scale of this stage is 1.5 ± 0.3 *µ*s. (B) State-B: The key molecular interaction induction stage of GDP extraction process. Four functional residues of SOS1 (Lys811, Glu812, Lys939 and Glu942) induce GDP to disengage from the GDP-binding pocket on KRAS, and the time scale of this stage is 2.9 ± 0.1 *µ*s. (C) Post-GDP extraction stage: GDP is released from KRAS and binds to the temporary GDP-binding pocket on the SOS1. SOS1 residues 930-944 are colored in “green surface”; SOS1 residues 807-818 are colored as an “ochre surface”. (D) The “Residue Cross Correlation” analysis of the KRAS^*WT*^ during the preparatory stage of GDP extraction. Highly (anti)correlated motions are depicted in deep blue(light yellow.) (E) The interaction model of the key molecular interactions during the “State-B”. Corresponding to (B) and all distances in Å. (F) The interaction mode of GDP binding to the temporary GDP-binding pocket on the SOS1 surface, corresponding to (C) and all distances in Å. (G) The SOS1-catalyzed GDP extraction process, characterized by “contact frequency” of the representative SOS1 residues with KRAS^*WT*^ and GDP.

As a complementary quantitative study of the interactions between different residues of SOS1 with those of GDP and KRAS, we are reporting a list of contact frequencies along the simulation time in Fig.2 G. The raw data of all independent trajectories are reported in the “Supplementary Information” (SI). Two particles/species are considered to be in “contact” when they form hydrogen bonds (HB) or salt bridges, with characteristic distances indicated in figures 2 E and F. Full details of the contact frequency are reported in Section4. The reported contact frequency map shows that the GDP and the guanine-free KRAS^*WT*^ tend to dissociate from the SOS1 surface. Considering that simulating the all-atom level KRAS-SOS1 interaction requires extremely huge computing resources and we have already observed the most important stages (“State-A” and “State-B”) of the GDP extraction catalyzed by SOS1, we stopped the simulation at 6 *µ*s (see Table1). As a summary of this part, we can state that: (1) KRAS^*WT*^, once loaded into the CDC25 catalytic site of SOS1, needs around 1.5 ± 0.3 *µ*s of KRAS^*WT*^ -SOS1 for an interface structural rearrangement, with *α*H opening/inserting into the SW-I region while IDRs^811−814^ approaches the P-loop and interacts with GDP; (2) subsequently, residues Lys811, Glu812, Lys939 and Glu942 are mainly involved in the establishment of the KRAS^*WT*^ -GDP-SOS1 molecular interaction induction interface and (3) after 2.9 ± 0.1 *µ*s of structural rearrangement, GDP is released from the guanosine binding site of KRAS^*WT*^ and it can be temporarily captured on the surface of SOS1 before being released to the solvent.

**Table 1.** The timescale statistics of different KRAS^*WT*^ -SOS1 interaction states.

| KRAS<sup>WT</sup>-SOS1 | Trajectory#1 | Trajectory#2 | Trajectory#3 | Average |
| --- | --- | --- | --- | --- |
| State-A | 1.6 μs | 1.2 μs | 1.6 μs | 1.5 ± 0.3 μs |
| State-B | 3.0 μs | 2.8 μs | 3.0 μs | 2.9 ± 0.1 μs |
| Total simulation time | 6.0 μs | 6.0 μs | 6.0 μs | - |

### 2.3 Molecular interactions of GDP extraction: KRAS-G12D case

Given that KRAS^*G*12*D*^ mutations are the most common and also the most elusive type of KRAS mutation-driven cancers, we have considered the molecular interactions of GDP extraction for the KRAS^*G*12*D*^ case. Basically, the main steps of GDP extraction by SOS1 from KRAS^*G*12*D*^-GDP complex are similar to those of the KRAS^*WT*^ case. The process of nucleotide extraction can be also divided into three consecutive steps. The obvious differences in comparison with the KRAS^*WT*^ case are as follows: (1) the SW-I domain of KRAS^*G*12*D*^ is significantly opened during the preparatory stage (Fig.3D); (2) KRAS^*G*12*D*^-SOS1 maintains a tight interaction interface (Fig.3G-State-A) and (3) the GDP extraction rate of KRAS^*G*12*D*^-SOS1 complex is significantly faster than that of the WT case.

**Figure 3.**
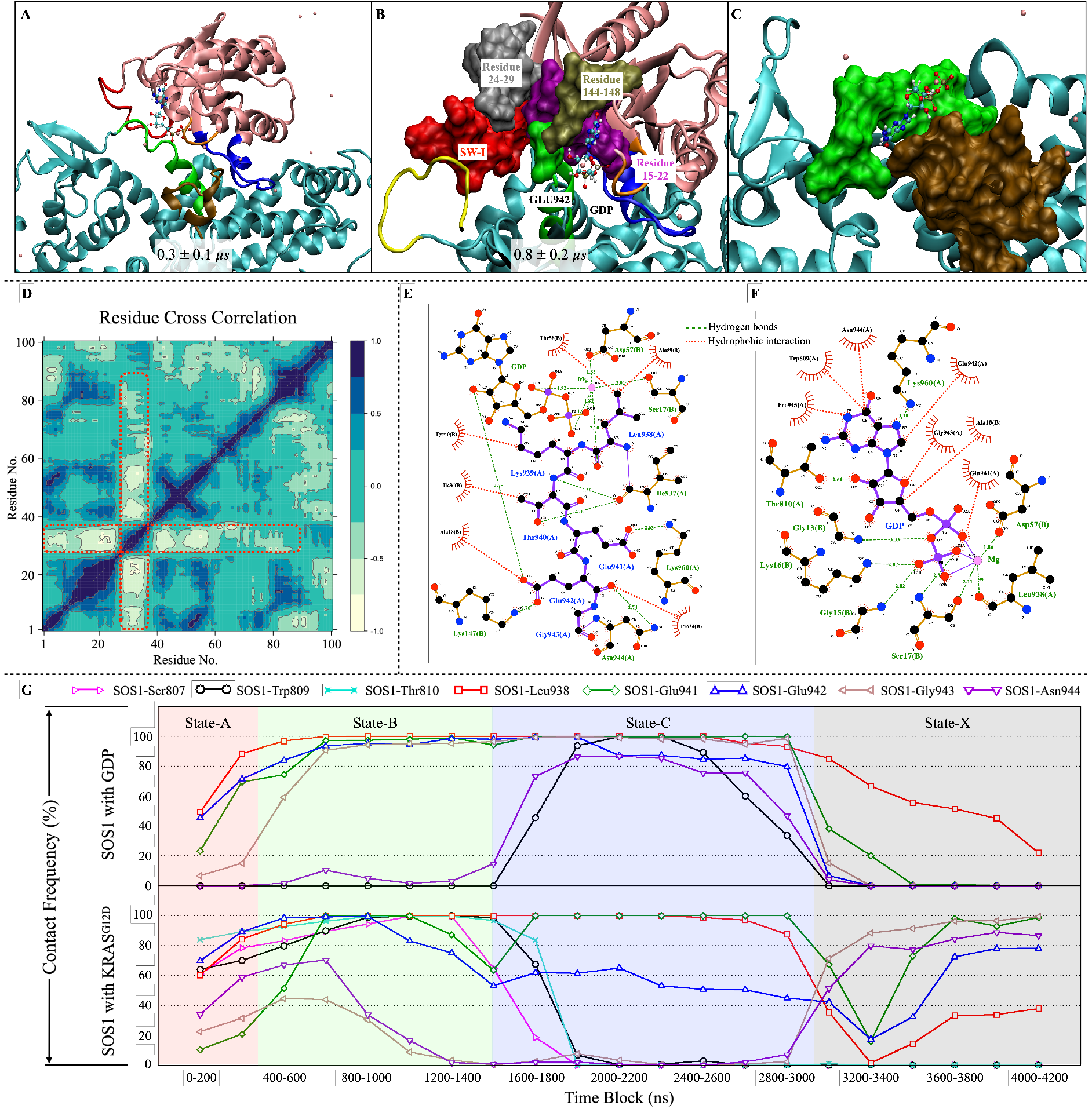
Molecular interactions of the GDP extraction process for the KRAS^*G*12*D*^-GDP-SOS1 complex. (A) State-A: Preparatory stage of GDP extraction. The schematic diagram of the KRAS^*G*12*D*^-SOS1 interactions in the preparation stage, and the time scale of this stage is 0.3 ± 0.1 *µ*s. (B) State-B: The key molecular interaction induction stage of GDP extraction process. The *α*H inserts deeper into the SW-I/GDP region and disrupts the GDP-binding pocket on KRAS^*G*12*D*^. The time scale of this stage is 0.8 ± 0.2 *µ*s. (C) Post-GDP extraction stage: GDP is released from KRAS^*G*12*D*^ and binds to the temporary GDP-binding pocket on the SOS1. SOS1 residues 930-944 is are colored as a “green surface”; SOS1 residues 807-818 are colored as an “ochre surface”. (D) The “Residue Cross Correlation” analysis of the KRAS^*G*12*D*^ during the preparatory stage of GDP extraction. Highly (anti)correlated motions are depicted in deep blue (light yellow). (E) The interaction model of the key molecular interactions during the “State-B”. Unlike the KRAS^*WT*^ case (Fig.2 E), the *α*H^938−943^ inserted into the SW-I region were taken as the main objects here. The residues (and GDP) that interact with them are distributed around the *α*H^938−943^. Corresponding to (B) and all distances in Å. (F) The interaction mode of GDP binding to the temporary GDP-binding pocket on the SOS1 surface, corresponding to (C) and all distances in Å. (G) The SOS1-catalyzed GDP extraction process, characterized by “contact frequency” of the representative SOS1 residues with KRAS^*G*12*D*^ and GDP.

The preparation process for GDP extraction (State-A) is approximately 4.5-fold faster than in the KRAS^*WT*^ case and the time required for the molecular interaction induction stage (State-B) is only about 30% of KRAS^*WT*^ ‘s one. Through the probability distribution of the distance between the 12th. amino acid (C*α*) of KRAS and the IDRs^807−818^ (geometric center of C*α*), it can be clearly seen that the proximity between the P-loop and the IDRs^807−818^ after KRAS-G12D mutation is the main reason for these changes (see Fig.4B). The significant reduction of the distance between the P-loop and the IDRs^807−818^ allows KRAS^*G*12*D*^-SOS1 to maintain a more compact interaction interface, making it easier for *α*H to access, open and insert deeper into the SW-I/GDP region, thereby shortening the “State-A” of GDP extraction. This is consistent with the “residue cross correlation” and the “contact frequency” trends in Fig.3 D and G. Moreover, the detailed atomic-level interaction model of Asp12 with IDRs^807−818^ can be understood with the help of Asp12-IDRs^807−818^ contact frequency: the negatively charged mutation Asp12 can undergo extensive hydrogen bonding interactions with polar residues on the IDRs^807−818^ region (see Fig.4A and Table3).

**Figure 4.**
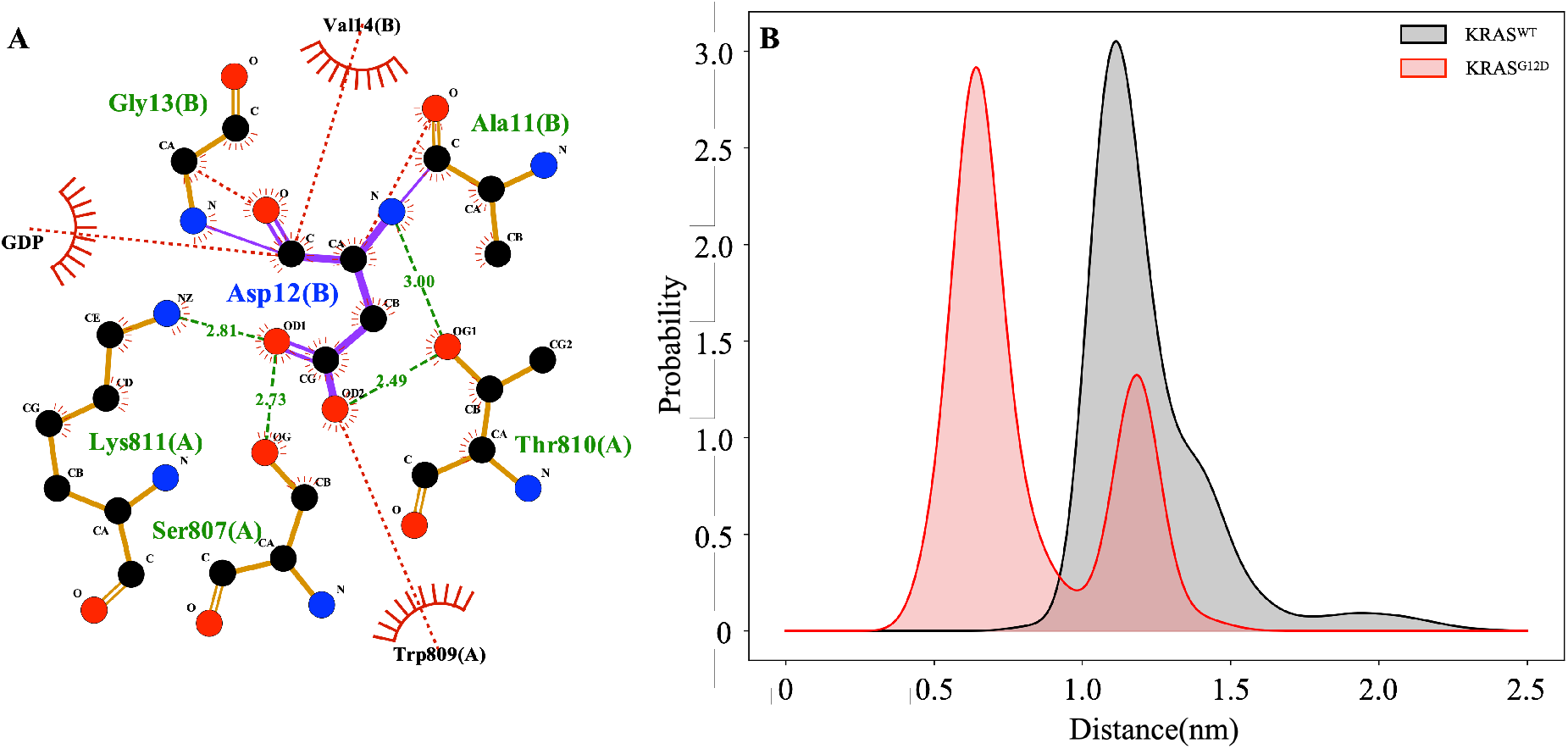
Corresponding atomistic mechanism of accelerated GDP release of the KRAS-G12D case. (A) The interaction model of KRAS(Asp12) with SOS1 (residues 804-819) during the preparatory stage of GDP extraction. (B) The probability distribution of the distance between the 12th. amino acid (C*α*) of KRAS and the SOS1 (C*α* of residues 807-818).

In the subsequent molecular interaction induction stage (State-B) the largely opened SW-I is stabilized by the IDRs^589−604^ domain of SOS1 (for more detailed imformation see the explicit contact tables at “Supplementary Information”). Furthermore, during the structural rearrangement of molecular interaction induction stage, *α*H^938−943^ can deconstruct the GDP-binding pocket of KRAS^*G*12*D*^ (see Fig.3B and E), ultimately leading to the extraction of GDP out of its bindig pocket at the KRAS surface. During the next post-GDP extraction stage, the KRAS^*G*12*D*^-SOS1 interaction interfaces are partly disconnected, such as the IDRs^807−818^ region, which in turn forms the temporary GDP-binding pocket with *α*H on the SOS1 surface so that GDP can be temporarily captured on this pocket, as described in Figs.3 C, F and G (labeld as State-C).

As the final step, the subsequent interaction interfaces of KRAS^*G*12*D*^-SOS1 are enhanced again and GDP is released from the SOS1 surface (see Fig.3 G, labeled as State-X). This suggests that the nucleotide-free RAS holds the high binding affinity with the SOS1 catalytic site, and this consistent well with previous experiments and theoretical calculations^46, 49, 50, 56^. Similarly to the KRAS^*WT*^ case, the simulation was stopped at 4.2 *µ*s and the other two replica simulations were stopped at 2.4 *µ*s (see Table2). The main conclusion of this Section is that the G12D mutation in KRAS greatly enhances the interaction between the P-loop and the IDRs^807−818^, making the KRAS^*G*12*D*^-SOS1 interaction interface much tighter than the KRAS^*WT*^ -SOS1. This produces two effects: (1) *α*F inserts deeper into the SW-I/GDP region and opens it largely; (2) The interaction of IDRs^807−818^ with the P-loop destabilizes the binding domain of the GDP phosphate group. The combination of these two factors accelerates the destruction of the GDP binding pocket and the release of GDP from KRAS^*G*12*D*^.

**Table 2.** Timescale statistics of different KRAS^*G*12*D*^-SOS1 interaction states.

| KRAS<sup>G12D</sup>-SOS1 | Trajectory#4 | Trajectory#5 | Trajectory#6 | Average |
| --- | --- | --- | --- | --- |
| State-A | 0.4 μs | 0.2 μs | 0.2 μs | 0.3 ± 0.1 μs |
| State-B | 1.0 μs | 0.8 μs | 0.6 μs | 0.8 ± 0.2 μs |
| Total simulation time | 4.2 μs | 2.4 μs | 2.4 μs | - |

**Table 3.**
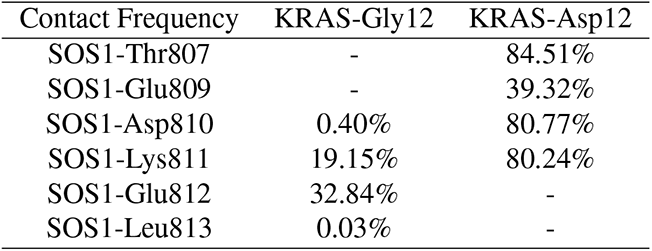
Contact frequencies between KRAS Gly12/Asp12 and SOS1 during the “State-A”.

| Contact Frequency | KRAS-Gly12 | KRAS-Asp12 |
| --- | --- | --- |
| SOS1-Thr807 | - | 84.51% |
| SOS1-Glu809 | - | 39.32% |
| SOS1-Asp810 | 0.40% | 80.77% |
| SOS1-Lys811 | 19.15% | 80.24% |
| SOS1-Glu812 | 32.84% | - |
| SOS1-Leu813 | 0.03% | - |

## 3. Discussion

Through microsecond scale all-atom MD simulations and trajectory analysis, the SOS1-catalyzed GDP extraction process and the underlying atomic-level molecular interactions are described. The IDRs^807−818^ and *α*H domains of SOS1 cooperatively participate in the GDP extraction process: *α*H opens and inserts into the SW-I domain of KRAS^*WT*^, while the IDRs^807−818^ interact with the phosphate group of GDP. Among them, residues Lys811, Glu812, Lys939 and Glu942 play the main role. Moreover, we also revealed the potential correlation between the enhanced GDP extraction rate and the KRAS-G12D mutation. Basically, the synergy of IDRs^807−818^ and *α*H is well-conserved during the GDP extraction steps and it is crucial for both KRAS^*WT*^ and KRAS^*G*12*D*^ cases. With the occurrence of G12D mutation, the molecular interactions at the atomic-level of GDP extraction slightly change and result in a significant increase of the GDP extraction rate. The negatively charged Asp12 mutation changes the charge distribution of the “non-polar” P-loop and greatly enhances the interactions with IDRs^807−818^, especially with residues Ser807, Trp809, Thr810 and Lys811. This destabilizes the phosphate binding loop and enables KRAS^*G*12*D*^-SOS1 to maintain a more compact interaction interface. Under the influence of the above-mentioned dual factors, the destruction of the nucleotide-binding pocket is accelerated, thereby promoting the release of GDP. Interestingly, when KRAS-G12C mutation occurs (the corresponding three independent trajectories are not reported here), GDP extraction speed is also accelerated taking only a total time of about 0.7 *µ*s, leading to the highest extraction rate of all three cases (WT, G12D and G12C).

The structure of the interaction interface of KRAS-SOS1 is critical to uncover the mechanism of the GDP extraction process. Detailed data on “contact frequencies” used to measure and monitor the structure of KRAS-SOS1 interface during GDP extraction are provided in the “Supplementary Information”. The KRAS^*WT*^ -SOS1 contact frequency shows that the main SOS1 residues involved in the KRAS^*WT*^ -SOS1 interaction interface are residues labeled 811-814, 828-836, 872-884, 905-913, 929, 936-944, 1003-1010 and 1019. The major SOS1 residues involved in the KRAS^*G*12*D*^-SOS1 interaction interface are similar to the KRAS^*WT*^ -SOS1 interface. The above KRAS-SOS1 interaction interface are in good agreement with previously published work^50^, indicating the reliability of the initial parameters of KRAS-SOS1 interface and the simulation results. The increased contact frequency of the IDRs^589−604^ domain in the KRAS^*G*12*D*^-SOS1 complex is mainly due to the fact that IDRs^589−604^ is involved in the stabilization of the largely opened SW-I of KRAS^*G*12*D*^.

To further compare with previous experimental results, the extracted KRAS-SOS1 interaction interface located at the boundary between “State-B” and post-GDP extraction stage was employed for superimposition comparison with existing crystal structures. The crystal structures employed are 1nvu, 1nvx and 1xd2, with the CDC25 catalytic site binds nucleotide-free HRAS (see Fig.5). Structural comparison shows that the structure of the CDC25 catalytic site after the GDP extraction process is well consistent with the existing crystal structure, while the IDRs^807−818^ domain is slightly offset. This provides strong support for the GDP extraction mechanisms based on the CDC25 catalytic site proposed in the present work. The slight shift of the IDRs^807−818^ domain is reasonable since: (1) the IDRs^807−818^ holds higher conformational freedom and (2) IDRs^807−818^ together with *α*H can also form a temporary GDP-binding pocket on the SOS1 surface.

**Figure 5.**
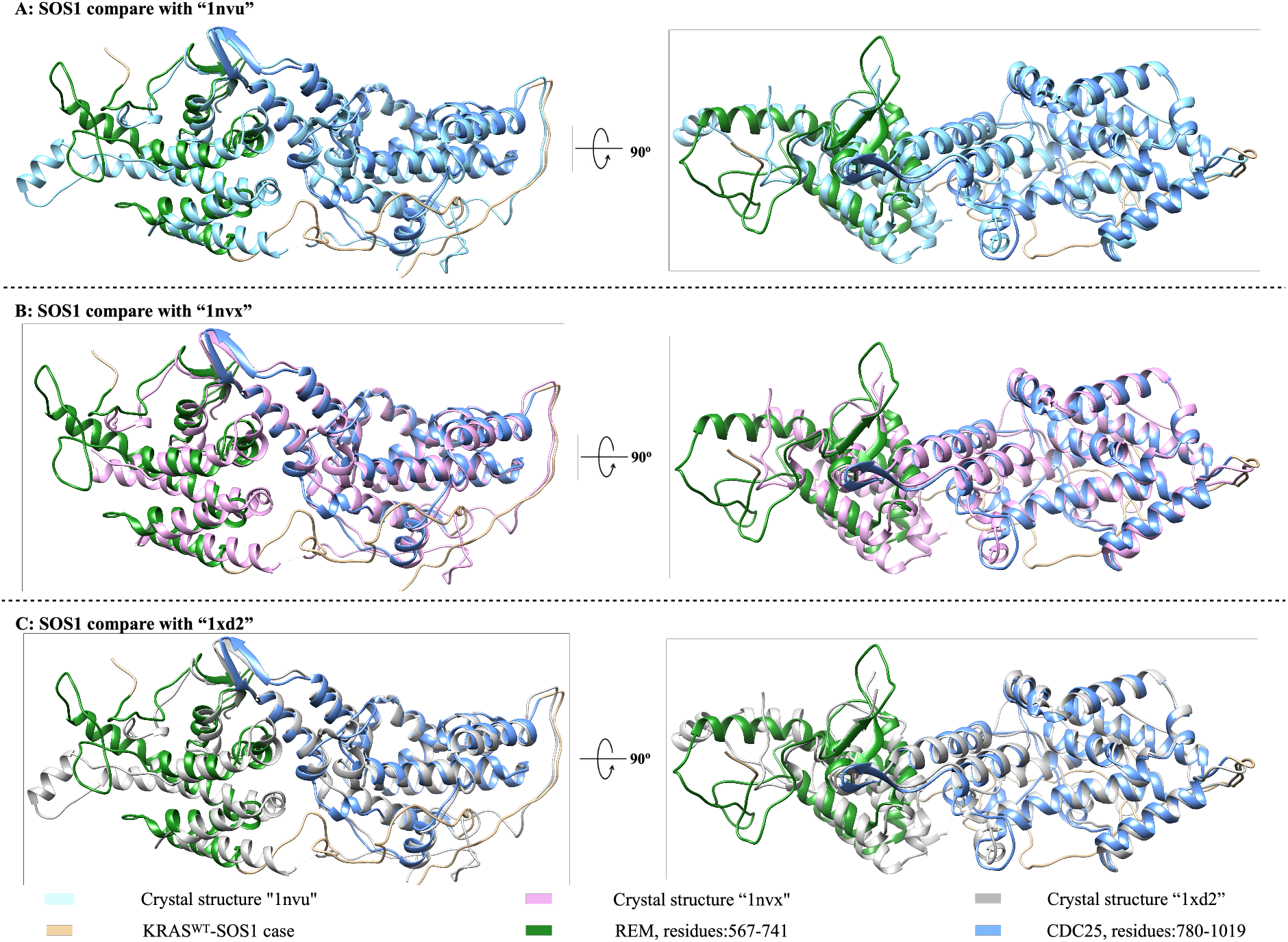
Experimental validation: superimposition of the SOS1 conformation and the relevant crystal structure after the releasing of GDP. (A) Superimposition of SOS1 conformation (KRAS-WT case) and crystal structure “1nvu”^45^; (B) Superimposition of SOS1 conformation (KRAS-WT case) and crystal structure “1nvx”^45^; (C) Superimposition of SOS1 conformation (KRAS-WT case) and crystal structure “1xd2”^46^.

In conclusion, we have described molecular interactions of the SOS1-catalyzed GDP extraction in full details at the atomic level, as well as the impact of the KRAS-G12D mutation on GDP extraction. This will help to extend the knowledge of SOS1 function and the detailed impact of KRAS-G12D mutation. The findings also suggest that the pocket identified on the IDRs^807−818^ region can be considered as a potential target for the KRAS^*G*12*D*^-SOS1 specific development of therapeutic strategies, following one of the most relevant strategies for cancer treatments, namely targeting the process of activation of KRAS^*G*12*D*^, in a similar fashion as reported by Hoffmann et al.^40^.

## 4 Methods

### 4.1 The initial configurations of Ras-SOS1 interface

The crystal structures of SOS1 (PDB: 1bkd)^50^ and GDP-bound KRAS (PDB: 4obe)^57^ were obtained from the Protein Data Bank^58^. As for SOS1 structure (1bkd), the SOS1 catalytic site is occupied by GDP/GTP-free HRas. Chimera’s “Model/Refine Loops” modeller^59^ was used to complete the missing amino acid in the SOS1 crystal structure. After removing the GDP/GTP-free HRas, the KRAS was docked to the surface of SOS1, generating the coordinates for the KRAS-SOS1 complex. Docking results show that KRAS preferentially binds to the SOS1 catalytic site, which is consistent with previous work^45, 46, 49, 50^ and indicates that the docking results are credible. Moreover the KRAS-SOS1 protein-protein interaction docking was performed by ClusPro server^51–54^. The “completed” structure of SOS1 (residue:568-1044) was used as “receptor”, the wild-type KRAS and KRAS-G12D were used as “ligand”. The construction method of KRAS-G12D mutant is the same as that of previously works^22^.

### 4.2 Molecular dynamics simulations parameters

Our main computational tools have been microsecond scale MD, which is well suited to model a wide variety of biosystems at all-atom level^60–63^. In the present work we mainly conducted 6 independent MD simulations of KRAS-SOS1 interactions and the initial configurations were subject to MD simulations in an aqueous environment. Each system contained one GDP bound KRAS-SOS1 complex fully solvated by 82,127 TIP3P water molecules^64^ in potassium chloride solution(0.15 M) and magnesium chloride(0.03 M) yielding a system size of around 260,000 atoms. All MD inputs were generated by means of the CHARMM-GUI solution builder^65–67^ assuming the CHARMM36m force field^68^. The different evolutions of his force field have been tested with high reliability throughout the last 20 years^68–70^. All bonds involving hydrogens were set to fixed lengths, allowing fluctuations of bond distances and angles for the remaining atoms.

All the KRAS-SOS1 interaction systems were energy minimised and well equilibrated (NVT ensemble) before generating the production MD. 6 independent production runs were performed within the NPT ensemble. The pressure and temperature were set at 1 atm and 310.15 K respectively. Pressure was controlled by a Parrinello-Rahman piston with damping coefficient of 5 ps^−1^ whereas temperature was controlled by a Nosé-Hoover thermostat with a damping coefficient of 1 ps^−1^. A time steps of 2 fs was used in all equilibration and production simulations and the particle mesh Ewald method with Coulomb radius of 1.2 nm was employed to compute long-ranged electrostatic interactions. The cutoff for Lennard-Jones interactions was set to 1.2 nm. Periodic boundary conditions in three directions of space have been taken.

### 4.3 Trajectory analysis and visualisation

The GROMACS/2021 package^71^ was employed for the MD simulations, the software VMD^72^, UCSF Chimera^73^ and LigPlot^+74^ for analysis and visualization purposes. Moreover, in order to better analyze, display and divide the above-mentioned GDP extraction process of SOS1 from the atomic scale, we also introduced statistical methods called “residue-residue contact frequency” (abbreviated as AB as “contact frequency”) to analyze the simulation results. “Contact frequency” is defined by the distance between protein amino acid residues (or between ligand molecules). When the distance between the two is less than 4 Å, then it is defined that they are in contact with each other. We calculated and evaluated the KRAS-SOS1 and GDP-SOS1 “contact frequency” throughout the entire process of SOS1 extracting GDP for both wild-type and G12D case. For curious, see the detailed “Supplementary Information”. All meaningful properties were averaged from 3 independent trajectories.

## 5 Acknowledgements

We thank financial support provided by the Spanish Ministry of Science, Innovation and Universities. This publication is a part of the I+D+i project with reference PID2021-124297NB-C32, founded by MCIN/AEI/10.13-039/501100011033 and “FEDER Una manera de hacer Europa”. Zheyao Hu is a Ph.D. fellow from the China Scholarship Council (grant 202006230070). J.M. thanks the *Generalitat de Catalunya* for the support through the grant *Grup de Recerca SGR-Cat2021 Condensed, Complex and Quantum Matter Group* reference 2021SGR-01411 and to the Polytechnic University of Catalonia-Barcelona Tech through the funding AGRUPS. Finally, computational resources provided by the Barcelona Supercomputing Center, project BCV-2023-2-0004 are also acknowledged.

## 6 Supporting Information

Full computational details are reported on “Supplementary Information”. The system composition, molecular dynamics simulation setups for the KRAS-SOS1 system can be seen on the Github: https://github.com/Zheyao-Hu/KRAS-SOS1

## 7 Addendum

Competing Interests: The authors declare that they have no competing financial interests.

