## Supplementary Information for "Atomic-scale mechanisms of GDP extraction by SOS1 in KRAS-G12 and KRAS-D12 oncogenes"

### ABSTRACT

The main mechanisms of GDP extraction from the KRAS oncogenes by means of the guanine exchange factor SOS1 are disclosed and described at the atomic-level. All contact frequency data used in the main text are listed in Tables 1-12.

### 1 Contact frequency statistics

**Time-dependent contact frequency statistics: frequency of SOS1 residues contacting KRAS-WT during the simulation (trajectory#1)**

| | State-A | | | | State-B | | | | | | | GDP release | | | | |
|---|---|---|---|---|---|---|---|---|---|---|---|---|---|---|---|---|
| **Residue** | **0-400** | **400-800** | **800-1200** | **1200-1600** | **1600-2000** | **2000-2400** | **2400-2800** | **2800-3200** | **3200-3600** | **3600-4000** | **4000-4400** | **4400-4800** | **4800-5200** | **5200-5600** | **5600-6000** | **……** |
| **SOS1-LYS-811** | 3.12% | 31.65% | 51.03% | 89.21% | 99.48% | 99.90% | 99.97% | 99.70% | 99.93% | 99.90% | 99.96% | 99.97% | 100.00% | 99.99% | 99.05% | |
| **SOS1-GLU-812** | 11.14% | 25.58% | 54.35% | 87.14% | 92.51% | 94.72% | 94.95% | 96.48% | 95.96% | 93.08% | 96.73% | 65.63% | 62.49% | 45.91% | 31.30% | |
| **SOS1-LYS-814** | 18.87% | 46.52% | 67.96% | 81.33% | 91.23% | 86.89% | 89.24% | 82.93% | 83.47% | 73.27% | 87.50% | 25.52% | 12.50% | 10.19% | 6.44% | |
| **SOS1-ILE-825** | 5.33% | 16.87% | 1.37% | 31.62% | 28.01% | 21.62% | 6.35% | 7.41% | 29.49% | 60.02% | 82.55% | 41.66% | 35.81% | 32.78% | 32.87% | |
| **SOS1-THR-828** | 29.29% | 31.05% | 54.11% | 80.80% | 95.73% | 96.66% | 89.28% | 87.74% | 90.53% | 97.97% | 97.14% | 96.59% | 96.80% | 96.65% | 97.82% | |
| **SOS1-THR-829** | 33.94% | 38.54% | 0.38% | 6.28% | 24.37% | 60.80% | 45.96% | 56.14% | 51.88% | 65.46% | 88.89% | 86.38% | 85.58% | 81.48% | 78.72% | |
| **SOS1-THR-832** | 33.83% | 73.43% | 14.80% | 33.63% | 64.86% | 99.73% | 91.16% | 96.83% | 96.78% | 99.99% | 99.97% | 99.82% | 99.90% | 99.75% | 99.93% | |
| **SOS1-GLU-836** | 43.22% | 51.63% | 81.95% | 89.26% | 99.57% | 99.91% | 94.41% | 100.00% | 100.00% | 100.00% | 100.00% | 100.00% | 99.82% | 100.00% | 100.00% | |
| **SOS1-LEU-872** | 71.13% | 87.39% | 93.50% | 92.09% | 92.23% | 95.68% | 96.23% | 91.09% | 92.21% | 98.42% | 96.87% | 88.71% | 90.37% | 90.42% | 90.87% | |
| **SOS1-GLU-873** | 31.01% | 41.64% | 55.84% | 60.49% | 93.83% | 99.26% | 98.53% | 99.16% | 98.82% | 99.43% | 98.82% | 91.47% | 94.10% | 96.24% | 96.15% | |
| **SOS1-VAL-875** | 71.29% | 81.37% | 96.90% | 93.35% | 85.78% | 85.77% | 94.65% | 92.76% | 90.58% | 81.32% | 84.36% | 78.88% | 83.92% | 80.20% | 85.74% | |
| **SOS1-SER-876** | 96.64% | 100.00% | 99.97% | 100.00% | 100.00% | 100.00% | 100.00% | 100.00% | 100.00% | 100.00% | 100.00% | 100.00% | 100.00% | 100.00% | 100.00% | |
| **SOS1-ASN-879** | 95.91% | 100.00% | 100.00% | 100.00% | 100.00% | 100.00% | 100.00% | 100.00% | 100.00% | 100.00% | 100.00% | 100.00% | 99.98% | 100.00% | 100.00% | |
| **SOS1-SER-880** | 94.20% | 99.76% | 97.79% | 99.66% | 99.97% | 99.95% | 99.84% | 99.90% | 99.93% | 99.86% | 99.85% | 99.90% | 99.95% | 99.99% | 99.99% | |
| **SOS1-SER-881** | 89.82% | 99.98% | 99.76% | 99.84% | 100.00% | 99.96% | 100.00% | 99.96% | 100.00% | 100.00% | 100.00% | 99.99% | 100.00% | 100.00% | 99.75% | |
| **SOS1-TYR-884** | 90.16% | 99.32% | 99.08% | 99.53% | 98.59% | 98.43% | 98.65% | 99.85% | 99.12% | 100.00% | 100.00% | 100.00% | 99.99% | 97.80% | 98.77% | |
| **SOS1-HIS-905** | 28.20% | 33.34% | 79.15% | 67.38% | 52.53% | 71.88% | 70.77% | 28.45% | 24.19% | 38.13% | 32.36% | 52.48% | 56.82% | 12.91% | 48.81% | |
| **SOS1-SER-908** | 83.40% | 81.15% | 83.65% | 87.15% | 87.72% | 76.27% | 89.46% | 90.37% | 86.61% | 88.28% | 82.59% | 93.91% | 89.86% | 83.77% | 93.65% | |
| **SOS1-GLU-909** | 36.37% | 58.95% | 78.64% | 69.81% | 61.03% | 70.07% | 76.02% | 73.44% | 59.91% | 55.46% | 62.14% | 63.84% | 62.49% | 57.16% | 67.52% | |
| **SOS1-ASP-910** | 27.42% | 13.30% | 8.40% | 54.91% | 65.12% | 49.79% | 62.83% | 62.80% | 53.47% | 40.06% | 38.14% | 51.91% | 76.03% | 47.08% | 69.86% | |
| **SOS1-HSD-911** | 59.50% | 69.25% | 99.95% | 99.89% | 99.97% | 99.99% | 100.00% | 100.00% | 99.93% | 100.00% | 99.93% | 99.15% | 99.70% | 99.91% | 98.60% | |
| **SOS1-TYR-912** | 86.56% | 91.43% | 89.70% | 94.32% | 93.77% | 96.05% | 96.32% | 97.46% | 97.74% | 97.30% | 98.42% | 92.96% | 93.58% | 87.50% | 87.15% | |
| **SOS1-LYS-913** | 20.75% | 31.70% | 81.05% | 96.68% | 94.03% | 94.27% | 92.24% | 97.37% | 94.49% | 92.74% | 95.33% | 94.20% | 92.88% | 93.61% | 87.37% | |
| **SOS1-PHE-929** | 54.91% | 77.63% | 79.42% | 83.45% | 69.79% | 86.67% | 86.81% | 64.64% | 78.60% | 94.41% | 94.25% | 95.01% | 84.45% | 78.34% | 90.92% | |
| **SOS1-ILE-932** | 9.89% | 7.64% | 26.17% | 39.45% | 43.72% | 17.62% | 9.70% | 0.34% | 0.00% | 0.00% | 0.00% | 0.26% | 0.04% | 5.15% | 0.23% | |
| **SOS1-ASN-936** | 3.75% | 24.42% | 45.74% | 70.60% | 71.97% | 78.80% | 86.67% | 96.22% | 96.44% | 93.48% | 76.63% | 74.04% | 39.52% | 39.12% | 20.60% | |
| **SOS1-LYS-939** | 15.65% | 73.73% | 85.85% | 92.26% | 99.67% | 99.59% | 99.44% | 99.31% | 100.00% | 99.80% | 99.79% | 99.43% | 99.00% | 93.18% | 98.09% | |
| **SOS1-THR-940** | 0.38% | 5.24% | 10.76% | 38.58% | 44.37% | 48.54% | 44.34% | 63.51% | 65.18% | 84.17% | 72.62% | 79.16% | 25.99% | 45.07% | 29.82% | |
| **SOS1-GLU-942** | 13.24% | 42.68% | 50.57% | 79.98% | 93.74% | 92.25% | 93.01% | 93.01% | 97.98% | 94.44% | 90.59% | 89.98% | 51.12% | 39.65% | 20.04% | |
| **SOS1-GLY-943** | 10.23% | 47.85% | 58.56% | 80.42% | 89.01% | 97.38% | 93.29% | 98.36% | 99.55% | 99.40% | 99.63% | 89.78% | 84.29% | 64.80% | 49.08% | |
| **SOS1-ASN-944** | 14.68% | 32.76% | 41.03% | 61.03% | 78.63% | 81.90% | 80.33% | 80.97% | 75.42% | 97.56% | 99.86% | 77.83% | 54.20% | 34.19% | 20.76% | |
| **SOS1-PRO-945** | 30.22% | 19.78% | 0.61% | 29.38% | 28.48% | 72.11% | 60.34% | 58.36% | 62.38% | 82.20% | 69.79% | 37.13% | 40.30% | 62.52% | 41.01% | |
| **SOS1-GLU-1002** | 11.34% | 16.40% | 7.28% | 5.68% | 3.73% | 11.42% | 5.85% | 8.06% | 11.51% | 6.62% | 7.45% | 7.24% | 5.59% | 8.08% | 2.97% | |
| **SOS1-LYS-1003** | 63.28% | 69.21% | 61.90% | 75.23% | 83.86% | 98.02% | 87.50% | 89.42% | 89.45% | 87.16% | 83.23% | 86.61% | 91.23% | 93.42% | 98.35% | |
| **SOS1-THR-1006** | 45.97% | 34.61% | 24.92% | 29.52% | 22.23% | 35.49% | 46.37% | 50.10% | 39.57% | 15.11% | 20.99% | 16.56% | 18.69% | 41.85% | 20.00% | |
| **SOS1-ASP-1007** | 62.38% | 96.35% | 96.68% | 95.47% | 98.83% | 99.99% | 99.14% | 98.46% | 97.54% | 95.70% | 96.29% | 86.05% | 79.46% | 79.94% | 69.99% | |
| **SOS1-PHE-1010** | 62.44% | 98.83% | 98.94% | 94.22% | 98.31% | 97.98% | 98.09% | 97.57% | 97.44% | 97.33% | 97.90% | 87.83% | 77.83% | 67.98% | 66.99% | |
| **SOS1-ARG-1019** | 44.01% | 87.07% | 87.99% | 73.27% | 75.18% | 44.21% | 75.16% | 71.81% | 70.00% | 69.49% | 74.31% | 62.03% | 59.41% | 42.29% | 39.15% | |
| | PS: Some residues with low contact frequency were not counted | | | | | | | | | | | | | | | |

**Figure 1.** Time-dependent contact frequency statistics: SOS1 residues contacting KRAS-WT during the simulation (trajectory#1).

**Time-dependent contact frequency statistics: frequency of SOS1 residues contacting GDP (KRAS-WT case) during the simulation (trajectory#1)**

| | State-A | | | | State-B | | | | | | | GDP release | | | |
|---|---|---|---|---|---|---|---|---|---|---|---|---|---|---|---|
| **Residue** | **0-400** | **400-800** | **800-1200** | **1200-1600** | **1600-2000** | **2000-2400** | **2400-2800** | **2800-3200** | **3200-3600** | **3600-4000** | **4000-4400** | **4400-4800** | **4800-5200** | **5200-5600** | **5600-6000** |
| **PROA-LYS-811** | 0.00% | 1.04% | 34.82% | 88.79% | 99.87% | 99.46% | 100.00% | 93.38% | 94.90% | 99.31% | 99.86% | 96.36% | 99.71% | 100.00% | 99.56% |
| **PROA-GLU-812** | 0.00% | 4.77% | 12.11% | 23.61% | 39.21% | 54.84% | 67.46% | 67.71% | 73.40% | 78.78% | 74.30% | 76.55% | 75.05% | 62.18% | 19.42% |
| **PROA-LYS-939** | 0.12% | 2.37% | 7.67% | 43.48% | 52.85% | 88.74% | 95.18% | 99.93% | 99.86% | 98.51% | 88.31% | 99.54% | 99.98% | 99.84% | 99.97% |
| **PROA-GLU-942** | 6.83% | 11.35% | 23.91% | 67.67% | 82.13% | 96.55% | 91.93% | 97.38% | 99.77% | 98.80% | 94.00% | 85.80% | 95.59% | 87.07% | 94.49% |
| | PS: Some residues with low contact frequency were not counted | | | | | | | | | | | | | | |

**Figure 2.** Time-dependent contact frequency statistics: SOS1 residues contacting GDP (KRAS-WT case) during the simulation (trajectory#1).

**Time-dependent contact frequency statistics: frequency of SOS1 residues contacting KRAS-WT during the simulation (trajectory#2)**

| | State-A | | | State-B | | | | | | | GDP release | | | | | |
|---|---|---|---|---|---|---|---|---|---|---|---|---|---|---|---|---|
| Residue | 0-400 | 400-800 | 800-1200 | 1200-1600 | 1600-2000 | 2000-2400 | 2400-2800 | 2800-3200 | 3200-3600 | 3600-4000 | 4000-4400 | 4400-4800 | 4800-5200 | 5200-5600 | 5600-6000 | ...... |
| SOS1-LYS-811 | 27.06% | 50.79% | 78.98% | 93.52% | 99.88% | 100.00% | 98.55% | 99.73% | 99.02% | 97.69% | 100.00% | 99.97% | 98.43% | 97.58% | 99.42% | |
| SOS1-GLU-812 | 33.02% | 46.63% | 79.62% | 90.46% | 94.61% | 95.36% | 96.05% | 98.96% | 99.73% | 95.58% | 94.62% | 70.03% | 54.89% | 39.88% | 28.32% | |
| SOS1-LYS-814 | 40.98% | 65.78% | 87.69% | 91.43% | 95.76% | 89.77% | 90.63% | 80.84% | 85.56% | 76.78% | 90.04% | 33.48% | 20.64% | 5.48% | 8..96% | |
| SOS1-THR-828 | 55.08% | 71.23% | 77.62% | 88.88% | 94.80% | 97.88% | 92.46% | 85.58% | 90.90% | 94.86% | 98.43% | 99.72% | 95.47% | 97.38% | 94.96% | |
| SOS1-THR-829 | 55.78% | 72.73% | 25.46% | 30.52% | 40.96% | 62.71% | 50.06% | 53.68% | 55.95% | 63.25% | 90.01% | 84.76% | 80.50% | 73.24% | 66.82% | |
| SOS1-THR-832 | 57.62% | 80.07% | 34.88% | 45.04% | 70.55% | 100.00% | 92.72% | 95.69% | 98.89% | 100.00% | 96.34% | 95.72% | 99.23% | 96.35% | 98.62% | |
| SOS1-GLU-836 | 60.02% | 82.71% | 90.03% | 91.28% | 95.88% | 97.69% | 100.00% | 99.32% | 97.24% | 99.89% | 100.00% | 96.58% | 98.32% | 99.66% | 98.87% | |
| SOS1-LEU-872 | 85.21% | 93.65% | 94.72% | 93.44% | 92.86% | 96.52% | 97.03% | 92.34% | 94.53% | 99.98% | 97.04% | 90.64% | 90.58% | 88.36% | 91.55% | |
| SOS1-GLU-873 | 50.20% | 63.74% | 75.62% | 82.41% | 94.34% | 100.00% | 97.22% | 99.01% | 97.60% | 98.82% | 99.35% | 90.26% | 95.18% | 94.38% | 97.21% | |
| SOS1-VAL-875 | 76.48% | 86.49% | 97.49% | 95.42% | 87.63% | 84.88% | 97.87% | 93.08% | 91.22% | 80.46% | 86.79% | 76.93% | 85.74% | 77.98% | 83.62% | |
| SOS1-SER-876 | 94.82% | 98.66% | 97.82% | 100.00% | 99.65% | 98.42% | 99.10% | 100.00% | 100.00% | 97.98% | 100.00% | 99.53% | 100.00% | 97.34% | 100.00% | |
| SOS1-ASN-879 | 96.08% | 99.30% | 99.78% | 100.00% | 100.00% | 97.84% | 99.76% | 100.00% | 98.40% | 100.00% | 99.15% | 98.88% | 99.65% | 100.00% | 99.02% | |
| SOS1-SER-880 | 88.32% | 93.80% | 95.92% | 100.00% | 99.52% | 99.04% | 95.79% | 99.46% | 100.00% | 99.78% | 98.68% | 98.60% | 99.44% | 97.84% | 100.00% | |
| SOS1-SER-881 | 87.64% | 100.00% | 100.00% | 99.06% | 99.68% | 97.69% | 98.77% | 100.00% | 99.61% | 98.36% | 99.59% | 100.00% | 99.28% | 97.27% | 98.61% | |
| SOS1-TYR-884 | 89.15% | 99.27% | 97.40% | 100.00% | 97.68% | 95.37% | 98.04% | 100.00% | 98.72% | 98.26% | 99.79% | 98.63% | 100.00% | 96.45% | 95.39% | |
| SOS1-HIS-905 | 36.22% | 50.64% | 80.02% | 65.43% | 55.59% | 75.65% | 72.97% | 35.03% | 26.30% | 40.52% | 30.47% | 55.91% | 45.42% | 17.44% | 46.08% | |
| SOS1-SER-908 | 80.41% | 82.20% | 84.25% | 90.05% | 84.33% | 78.88% | 90.04% | 91.21% | 84.93% | 87.89% | 80.46% | 95.85% | 87.68% | 80.06% | 88.43% | |
| SOS1-GLU-909 | 44.42% | 60.78% | 80.32% | 75.77% | 63.46% | 68.88% | 78.70% | 76.04% | 60.73% | 56.58% | 60.60% | 66.76% | 59.43% | 50.87% | 56.15% | |
| SOS1-ASP-910 | 33.75% | 16.72% | 10.47% | 59.02% | 67.84% | 45.27% | 59.88% | 63.26% | 56.96% | 39.46% | 37.43% | 58.93% | 74.41% | 50.86% | 67.28% | |
| SOS1-HSD-911 | 54.53% | 71.55% | 100.00% | 100.00% | 97.45% | 95.23% | 97.54% | 98.89% | 99.72% | 99.47% | 100.00% | 97.25% | 99.37% | 98.56% | 95.63% | |
| SOS1-TYR-912 | 83.42% | 92.78% | 86.47% | 95.50% | 94.12% | 97.25% | 98.48% | 98.26% | 95.44% | 98.03% | 99.22% | 90.18% | 92.56% | 85.35% | 83.33% | |
| SOS1-LYS-913 | 39.86% | 52.27% | 84.15% | 98.73% | 95.60% | 91.97% | 94.63% | 98.66% | 95.34% | 89.94% | 93.92% | 96.26% | 95.55% | 90.42% | 85.64% | |
| SOS1-PHE-929 | 60.73% | 74.59% | 83.76% | 86.62% | 70.31% | 89.59% | 87.36% | 66.62% | 75.62% | 95.20% | 93.05% | 96.43% | 82.30% | 76.56% | 86.57% | |
| SOS1-ASN-936 | 20.83% | 44.68% | 50.59% | 68.62% | 73.24% | 77.58% | 90.04% | 97.35% | 97.62% | 92.30% | 80.78% | 75.24% | 45.16% | 36.23% | 26.77% | |
| SOS1-LYS-939 | 24.23% | 77.68% | 88.89% | 93.04% | 100.00% | 100.00% | 98.33% | 97.60% | 99.58% | 100.00% | 98.89% | 99.27% | 97.08% | 95.36% | 97.64% | |
| SOS1-THR-940 | 4.96% | 19.34% | 24.93% | 44.76% | 46.03% | 50.38% | 46.71% | 58.78% | 66.63% | 79.99% | 75.43% | 69.70% | 30.82% | 47.25% | 26.37% | |
| SOS1-GLU-942 | 32.11% | 53.42% | 65.92% | 83.28% | 95.40% | 90.78% | 94.13% | 97.04% | 98.61% | 94.30% | 89.39% | 84.58% | 53.28% | 33.25% | 16.34% | |
| SOS1-GLY-943 | 17.78% | 51.34% | 63.62% | 77.83% | 88.88% | 96.40% | 94.52% | 97.64% | 100.00% | 98.74% | 97.48% | 86.22% | 77.69% | 63.87% | 38.46% | |
| SOS1-ASN-944 | 12.24% | 40.30% | 47.87% | 66.86% | 75.69% | 83.49% | 82.78% | 83.28% | 79.02% | 95.31% | 100.00% | 79.89% | 60.33% | 40.76% | 21.03% | |
| SOS1-PRO-945 | 33.46% | 20.03% | 6.86% | 35.94% | 40.87% | 73.01% | 62.55% | 54.08% | 59.98% | 78.89% | 70.36% | 36.59% | 42.36% | 57.68% | 37.65% | |
| SOS1-LYS-1003 | 59.98% | 64.82% | 65.29% | 76.47% | 85.66% | 97.93% | 86.25% | 91.49% | 89.86% | 88.56% | 85.72% | 89.04% | 93.46% | 94.54% | 97.44% | |
| SOS1-THR-1006 | 40.44% | 36.78% | 22.88% | 30.43% | 24.67% | 36.97% | 44.66% | 52.21% | 40.63% | 13.14% | 22.88% | 17.92% | 20.73% | 39.43% | 19.68% | |
| SOS1-ASP-1007 | 58.57% | 94.31% | 97.33% | 93.69% | 98.46% | 100.00% | 98.76% | 96.86% | 98.63% | 94.36% | 97.54% | 84.21% | 80.55% | 76.43% | 64.71% | |
| SOS1-PHE-1010 | 59.72% | 95.96% | 97.99% | 96.44% | 98.65% | 95.43% | 98.24% | 96.61% | 98.70% | 95.76% | 96.48% | 85.26% | 74.23% | 59.79% | 57.64% | |
| SOS1-ARG-1019 | 46.40% | 84.93% | 89.78% | 75.73% | 76.79% | 50.46% | 80.05% | 75.66% | 73.20% | 66.75% | 70.87% | 67.45% | 55.35% | 39.97% | 34.70% | |
| | PS: Some residues with low contact frequency were not counted | | | | | | | | | | | | | | | |

**Figure 3.** Time-dependent contact frequency statistics: SOS1 residues contacting KRAS-WT during the simulation (trajectory#2).

**Time-dependent contact frequency statistics: frequency of SOS1 residues contacting GDP (KRAS-WT case) during the simulation (trajectory#2)**

| | State-A | | | State-B | | | | | | | GDP release | | | | |
|---|---|---|---|---|---|---|---|---|---|---|---|---|---|---|---|
| Residue | 0-400 | 400-800 | 800-1200 | 1200-1600 | 1600-2000 | 2000-2400 | 2400-2800 | 2800-3200 | 3200-3600 | 3600-4000 | 4000-4400 | 4400-4800 | 4800-5200 | 5200-5600 | 5600-6000 |
| PROA-LYS-811 | 13.78% | 25.46% | 77.04% | 93.23% | 100.00% | 97.88% | 99.72% | 95.46% | 95.02% | 97.43% | 100.00% | 94.74% | 98.62% | 97.89% | 95.42% |
| PROA-GLU-812 | 7.69% | 19.84% | 53.52% | 60.02% | 61.40% | 64.06% | 65.75% | 72.89% | 77.34% | 74.69% | 75.13% | 77.46% | 67.28% | 59.88% | 17.06% |
| PROA-LYS-939 | 10.82% | 21.60% | 48.78% | 79.36% | 80.03% | 89.49% | 94.68% | 100.00% | 98.73% | 99.62% | 87.43% | 98.46% | 100.00% | 97.73% | 98.88% |
| PROA-GLU-942 | 26.22% | 37.47% | 75.63% | 83.59% | 94.92% | 97.65% | 90.82% | 95.25% | 98.97% | 98.08% | 90.73% | 86.48% | 94.60% | 84.44% | 92.98% |
| | PS: Some residues with low contact frequency were not counted | | | | | | | | | | | | | | |

**Figure 4.** Time-dependent contact frequency statistics: SOS1 residues contacting GDP (KRAS-WT case) during the simulation (trajectory#2).

**Time-dependent contact frequency statistics: frequency of SOS1 residues contacting KRAS-WT during the simulation (trajectory#3)**

| | State-A | | | | State-B | | | | | | | GDP release | | | | |
|---|---|---|---|---|---|---|---|---|---|---|---|---|---|---|---|---|
| **Residue** | **0-400** | **400-800** | **800-1200** | **1200-1600** | **1600-2000** | **2000-2400** | **2400-2800** | **2800-3200** | **3200-3600** | **3600-4000** | **4000-4400** | **4400-4800** | **4800-5200** | **5200-5600** | **5600-6000** | **……** |
| **SOS1-LYS-811** | 4.18% | 29.98% | 50.62% | 87.28% | 97.61% | 98.29% | 99.06% | 97.72% | 98.04% | 100.00% | 97.60% | 98.31% | 99.42% | 99.22% | 98.12% | |
| **SOS1-GLU-812** | 9.74% | 23.69% | 53.74% | 85.23% | 90.42% | 95.48% | 93.25% | 95.88% | 96.46% | 94.24% | 94.89% | 66.04% | 60.37% | 47.02% | 34.32% | |
| **SOS1-LYS-814** | 15.67% | 44.72% | 70.02% | 79.84% | 92.34% | 85.89% | 87.62% | 84.44% | 85.86% | 74.48% | 86.51% | 30.82% | 21.05% | 12.08% | 10.76% | |
| **SOS1-ILE-825** | 4.02% | 17.98% | 4.78% | 30.02% | 30.20% | 22.74% | 4.22% | 6.12% | 30.58% | 57.92% | 80.72% | 43.76% | 34.42% | 33.64% | 31.34% | |
| **SOS1-THR-828** | 27.08% | 30.14% | 53.12% | 81.83% | 96.81% | 97.32% | 87.69% | 88.52% | 91.64% | 96.75% | 96.63% | 97.19% | 95.82% | 97.58% | 94.74% | |
| **SOS1-THR-829** | 30.52% | 36.84% | 2.52% | 10.02% | 23.45% | 62.83% | 50.14% | 57.26% | 49.76% | 63.85% | 87.98% | 87.40% | 88.69% | 82.02% | 80.02% | |
| **SOS1-THR-832** | 34.72% | 72.48% | 17.60% | 35.74% | 70.90% | 97.63% | 94.46% | 97.54% | 93.34% | 98.78% | 97.98% | 100.00% | 96.28% | 98.04% | 99.24% | |
| **SOS1-GLU-836** | 40.14% | 52.90% | 83.60% | 88.42% | 98.12% | 99.10% | 96.32% | 100.00% | 100.00% | 97.56% | 100.00% | 98.72% | 99.44% | 100.00% | 99.02% | |
| **SOS1-LEU-872** | 70.18% | 85.32% | 94.51% | 94.16% | 95.03% | 95.18% | 97.45% | 90.18% | 94.54% | 97.68% | 95.92% | 90.02% | 92.76% | 91.34% | 89.86% | |
| **SOS1-GLU-873** | 29.72% | 39.72% | 56.04% | 65.02% | 92.44% | 97.32% | 98.26% | 96.96% | 97.54% | 99.30% | 97.98% | 93.56% | 95.15% | 95.72% | 94.50% | |
| **SOS1-VAL-875** | 68.43% | 83.69% | 94.92% | 90.52% | 86.56% | 90.76% | 95.46% | 96.06% | 90.14% | 80.12% | 85.76% | 75.69% | 85.48% | 82.48% | 86.02% | |
| **SOS1-SER-876** | 90.43% | 99.02% | 100.00% | 98.36% | 99.24% | 100.00% | 99.09% | 100.00% | 100.00% | 97.63% | 100.00% | 94.88% | 100.00% | 100.00% | 100.00% | |
| **SOS1-ASN-879** | 92.12% | 96.86% | 100.00% | 100.00% | 100.00% | 100.00% | 100.00% | 100.00% | 98.32% | 100.00% | 100.00% | 100.00% | 99.98% | 100.00% | 98.80% | |
| **SOS1-SER-880** | 91.32% | 97.68% | 98.69% | 99.24% | 98.02% | 99.84% | 97.63% | 99.92% | 97.54% | 98.70% | 98.65% | 97.68% | 98.66% | 99.96% | 99.26% | |
| **SOS1-SER-881** | 80.52% | 95.25% | 96.78% | 98.98% | 100.00% | 98.08% | 99.73% | 99.92% | 100.00% | 99.12% | 98.04% | 99.78% | 100.00% | 97.63% | 98.28% | |
| **SOS1-TYR-884** | 85.76% | 97.64% | 97.16% | 96.06% | 99.08% | 97.63% | 95.48% | 98.88% | 97.24% | 100.00% | 99.54% | 100.00% | 98.97% | 98.80% | 96.86% | |
| **SOS1-HIS-905** | 30.21% | 37.84% | 80.55% | 65.46% | 56.74% | 74.68% | 71.68% | 30.15% | 26.48% | 36.04% | 30.56% | 50.22% | 57.92% | 14.28% | 50.71% | |
| **SOS1-SER-908** | 80.23% | 80.76% | 85.94% | 90.05% | 86.54% | 77.67% | 90.66% | 90.02% | 84.40% | 87.64% | 84.69% | 95.02% | 88.76% | 84.87% | 95.08% | |
| **SOS1-GLU-909** | 40.36% | 60.06% | 77.48% | 70.14% | 60.84% | 75.03% | 77.01% | 68.84% | 60.80% | 54.36% | 65.04% | 64.40% | 60.38% | 56.64% | 64.28% | |
| **SOS1-ASP-910** | 30.14% | 15.34% | 7.80% | 50.03% | 64.24% | 50.83% | 65.14% | 60.81% | 55.48% | 43.66% | 35.84% | 56.70% | 75.34% | 44.02% | 65.46% | |
| **SOS1-HSD-911** | 60.45% | 67.48% | 97.85% | 99.08% | 98.84% | 99.10% | 100.00% | 99.24% | 98.30% | 100.00% | 100.00% | 98.01% | 99.26% | 97.90% | 99.33% | |
| **SOS1-TYR-912** | 84.62% | 90.73% | 88.63% | 95.64% | 94.08% | 97.65% | 98.36% | 96.63% | 98.82% | 98.36% | 96.38% | 91.86% | 94.66% | 85.52% | 89.30% | |
| **SOS1-LYS-913** | 22.25% | 32.68% | 84.45% | 97.44% | 93.62% | 95.54% | 91.48% | 98.58% | 95.50% | 91.23% | 96.43% | 95.02% | 94.26% | 90.52% | 86.88% | |
| **SOS1-PHE-929** | 52.54% | 74.83% | 80.48% | 84.33% | 70.49% | 87.32% | 88.84% | 65.62% | 80.24% | 95.38% | 96.45% | 97.72% | 86.56% | 80.04% | 90.92% | |
| **SOS1-ILE-932** | 11.09% | 8.20% | 30.28% | 40.36% | 45.78% | 20.88% | 11.06% | 5.04% | 0.86% | 1.28% | 0.00% | 0.42% | 0.16% | 7.01% | 0.78% | |
| **SOS1-ASN-936** | 6.83% | 30.02% | 46.86% | 67.96% | 70.98% | 80.28% | 91.36% | 97.28% | 95.76% | 88.63% | 73.64% | 70.84% | 45.02% | 39.86% | 22.67% | |
| **SOS1-LYS-939** | 13.78% | 70.62% | 88.80% | 94.56% | 98.77% | 100.00% | 97.48% | 98.23% | 99.76% | 100.00% | 97.82% | 98.30% | 99.18% | 95.78% | 97.29% | |
| **SOS1-THR-940** | 2.46% | 7.49% | 17.88% | 35.98% | 43.27% | 50.55% | 40.78% | 60.22% | 70.08% | 85.66% | 75.68% | 80.04% | 30.98% | 44.57% | 31.68% | |
| **SOS1-GLU-942** | 16.44% | 40.38% | 53.56% | 82.78% | 94.44% | 91.05% | 94.21% | 95.20% | 98.86% | 93.64% | 91.60% | 85.58% | 54.33% | 36.68% | 22.44% | |
| **SOS1-GLY-943** | 15.63% | 44.65% | 60.26% | 77.46% | 90.06% | 98.40% | 94.48% | 99.62% | 98.05% | 98.46% | 97.43% | 87.64% | 85.30% | 60.88% | 50.26% | |
| **SOS1-ASN-944** | 12.37% | 34.86% | 45.40% | 65.56% | 77.69% | 82.38% | 81.66% | 83.40% | 77.82% | 95.66% | 100.00% | 81.62% | 60.32% | 37.49% | 23.82% | |
| **SOS1-PRO-945** | 27.82% | 21.88% | 3.02% | 30.78% | 34.36% | 75.23% | 62.44% | 56.62% | 67.46% | 84.32% | 70.58% | 35.32% | 42.31% | 60.48% | 39.18% | |
| **SOS1-GLU-1002** | 14.53% | 20.42% | 11.39% | 7.88% | 4.03% | 15.20% | 3.66% | 10.20% | 9.62% | 5.84% | 6.38% | 8.72% | 6.33% | 8.35% | 3.48% | |
| **SOS1-LYS-1003** | 60.46% | 71.40% | 60.59% | 76.48% | 86.74% | 100.00% | 90.02% | 92.02% | 88.23% | 89.72% | 80.45% | 88.52% | 93.33% | 90.37% | 99.46% | |
| **SOS1-THR-1006** | 50.86% | 37.58% | 23.81% | 30.56% | 25.02% | 36.54% | 44.86% | 52.24% | 35.60% | 16.23% | 23.48% | 14.78% | 20.74% | 45.04% | 22.30% | |
| **SOS1-ASP-1007** | 60.00% | 92.24% | 97.36% | 93.50% | 96.78% | 100.00% | 98.06% | 99.30% | 97.48% | 94.87% | 97.40% | 85.45% | 82.35% | 80.24% | 68.79% | |
| **SOS1-PHE-1010** | 65.80% | 96.52% | 97.84% | 95.46% | 99.02% | 97.28% | 98.32% | 96.35% | 98.54% | 96.42% | 96.39% | 88.89% | 75.74% | 70.46% | 67.69% | |
| **SOS1-ARG-1019** | 40.30% | 84.38% | 89.09% | 70.46% | 77.42% | 50.34% | 73.84% | 70.98% | 70.07% | 67.88% | 77.42% | 65.32% | 57.84% | 40.46% | 40.37% | |
| | PS: Some residues with low contact frequency were not counted | | | | | | | | | | | | | | | |

**Figure 5.** Time-dependent contact frequency statistics: SOS1 residues contacting KRAS-WT during the simulation (trajectory#3).

**Time-dependent contact frequency statistics: frequency of SOS1 residues contacting GDP (KRAS-WT case) during the simulation (trajectory#3)**

| | State-A | | | | State-B | | | | | | | GDP release | | | |
|---|---|---|---|---|---|---|---|---|---|---|---|---|---|---|---|
| **Residue** | **0-400** | **400-800** | **800-1200** | **1200-1600** | **1600-2000** | **2000-2400** | **2400-2800** | **2800-3200** | **3200-3600** | **3600-4000** | **4000-4400** | **4400-4800** | **4800-5200** | **5200-5600** | **5600-6000** |
| **PROA-LYS-811** | 0.00% | 0.33% | 30.31% | 87.82% | 98.12% | 98.98% | 100.00% | 94.76% | 95.29% | 98.24% | 96.96% | 95.02% | 98.60% | 99.02% | 97.93% |
| **PROA-GLU-812** | 0.00% | 3.07% | 11.12% | 22.58% | 40.71% | 53.64% | 65.96% | 66.83% | 71.94% | 76.98% | 70.37% | 73.62% | 74.76% | 60.28% | 23.02% |
| **PROA-LYS-939** | 0.04% | 3.04% | 10.36% | 40.98% | 54.74% | 87.44% | 93.08% | 98.73% | 97.98% | 99.64% | 86.48% | 97.94% | 99.01% | 97.88% | 98.84% |
| **PROA-GLU-942** | 4.05% | 12.78% | 24.82% | 65.72% | 80.08% | 95.64% | 92.02% | 94.64% | 97.69% | 98.14% | 95.73% | 84.32% | 90.98% | 88.68% | 95.63% |
| | PS: Some residues with low contact frequency were not counted | | | | | | | | | | | | | | |

**Figure 6.** Time-dependent contact frequency statistics: SOS1 residues contacting GDP (KRAS-WT case) during the simulation (trajectory#3).

**Time-dependent contact frequency statistics: frequency of SOS1 residues contacting KRAS-G12D during the simulation (trajectory#4)**

| | State-A | | State-B | | | | | GDP release | | | | | | | | | | | | | | |
|---|---|---|---|---|---|---|---|---|---|---|---|---|---|---|---|---|---|---|---|---|---|---|
| **Residue** | **0-200** | **200-400** | **400-600** | **600-800** | **800-1000** | **1000-1200** | **1200-1400** | **1400-1600** | **1600-1800** | **1800-2000** | **2000-2200** | **2200-2400** | **2400-2600** | **2600-2800** | **2800-3000** | **3000-3200** | **3200-3400** | **3400-3600** | **3600-3800** | **3800-4000** | **4000-4200** | **……** |
| **SOS1-GLU-589** | 0.00% | 25.95% | 11.62% | 24.17% | 37.31% | 29.93% | 52.72% | 6.74% | 0.82% | 0.04% | 0.00% | 0.06% | 0.00% | 0.72% | 12.93% | 5.70% | 5.96% | 5.96% | 4.34% | 0.00% | 0.24% | |
| **SOS1-GLU-590** | 0.14% | 9.96% | 35.97% | 44.10% | 11.18% | 5.36% | 2.82% | 0.86% | 0.92% | 0.08% | 0.00% | 0.00% | 0.00% | 9.60% | 0.20% | 0.42% | 3.64% | 0.18% | 0.16% | 0.00% | 0.02% | |
| **SOS1-ASN-591** | 0.06% | 6.60% | 51.42% | 35.29% | 4.56% | 11.80% | 47.22% | 29.55% | 18.95% | 2.36% | 0.20% | 0.34% | 0.00% | 0.48% | 2.92% | 0.24% | 34.79% | 15.03% | 0.30% | 0.00% | 0.24% | |
| **SOS1-MET-592** | 0.04% | 0.34% | 7.22% | 51.72% | 19.51% | 33.61% | 94.74% | 74.09% | 66.15% | 16.67% | 9.22% | 13.09% | 0.84% | 26.65% | 16.19% | 5.60% | 39.48% | 13.55% | 1.10% | 0.38% | 1.00% | |
| **SOS1-GLN-593** | 0.00% | 0.00% | 0.00% | 0.00% | 0.00% | 14.89% | 76.87% | 53.04% | 61.26% | 36.11% | 46.74% | 27.21% | 1.36% | 9.54% | 17.19% | 3.54% | 6.80% | 1.38% | 1.42% | 0.14% | 1.58% | |
| **SOS1-PRO-594** | 0.00% | 0.00% | 0.00% | 0.00% | 0.00% | 8.12% | 59.32% | 33.91% | 16.77% | 31.01% | 76.29% | 44.70% | 0.74% | 0.44% | 4.80% | 0.68% | 2.74% | 0.38% | 0.00% | 0.00% | 0.00% | |
| **SOS1-LYS-595** | 0.00% | 0.00% | 0.00% | 0.00% | 0.00% | 0.00% | 25.99% | 8.74% | 0.46% | 4.76% | 12.91% | 12.83% | 0.02% | 2.12% | 0.08% | 0.00% | 0.00% | 0.00% | 0.94% | 0.00% | 0.60% | |
| **SOS1-ILE-600** | 0.00% | 0.00% | 0.00% | 0.00% | 0.00% | 12.53% | 54.96% | 23.29% | 12.38% | 0.94% | 0.00% | 0.00% | 0.00% | 0.00% | 0.36% | 0.08% | 0.02% | 0.04% | 0.00% | 0.00% | 0.00% | |
| **SOS1-LYS-602** | 1.24% | 52.98% | 32.33% | 44.82% | 81.99% | 43.64% | 51.48% | 7.24% | 6.04% | 10.01% | 3.47% | 0.00% | 5.41% | 0.06% | 3.08% | 6.26% | 27.37% | 18.57% | 4.06% | 0.08% | 0.34% | |
| **SOS1-SER-807** | 61.20% | 78.51% | 83.29% | 89.45% | 94.41% | 99.54% | 99.70% | 64.89% | 18.67% | 0.00% | 0.00% | 0.00% | 0.00% | 0.00% | 0.00% | 0.00% | 0.00% | 0.00% | 0.00% | 0.00% | 0.00% | |
| **SOS1-VAL-808** | 0.12% | 5.70% | 5.58% | 22.81% | 27.92% | 25.55% | 28.46% | 23.75% | 19.35% | 0.08% | 0.72% | 0.56% | 0.13% | 0.00% | 0.00% | 0.00% | 0.00% | 0.00% | 0.00% | 0.00% | 0.00% | |
| **SOS1-TRP-809** | 63.99% | 70.11% | 79.82% | 89.94% | 98.88% | 99.50% | 100.00% | 98.56% | 67.51% | 6.76% | 0.24% | 0.72% | 2.94% | 0.00% | 0.00% | 0.00% | 0.00% | 0.00% | 0.00% | 0.00% | 0.00% | |
| **SOS1-THR-810** | 83.87% | 89.24% | 92.68% | 96.28% | 99.40% | 99.56% | 99.56% | 96.78% | 83.53% | 0.02% | 0.00% | 0.00% | 0.00% | 0.00% | 0.00% | 1.02% | 0.00% | 0.00% | 0.00% | 0.00% | 0.00% | |
| **SOS1-LYS-811** | 64.31% | 77.66% | 77.69% | 78.13% | 77.31% | 71.96% | 71.70% | 54.90% | 20.47% | 0.00% | 0.00% | 0.00% | 0.00% | 0.00% | 0.00% | 1.32% | 0.00% | 0.00% | 0.00% | 0.00% | 0.00% | |
| **SOS1-GLU-812** | 9.56% | 20.31% | 56.46% | 58.36% | 26.55% | 18.95% | 19.01% | 15.93% | 6.66% | 3.40% | 0.00% | 0.00% | 1.00% | 0.22% | 0.52% | 2.58% | 0.37% | 0.77% | 0.00% | 3.80% | 9.36% | |
| **SOS1-LYS-814** | 62.36% | 74.13% | 86.35% | 95.36% | 63.32% | 49.45% | 42.76% | 22.72% | 10.84% | 3.24% | 0.00% | 0.00% | 0.36% | 0.14% | 5.62% | 8.32% | 10.20% | 24.19% | 24.57% | 39.16% | 73.10% | |
| **SOS1-SER-876** | 78.89% | 89.50% | 95.76% | 99.46% | 97.96% | 90.74% | 93.18% | 94.90% | 93.40% | 97.70% | 95.16% | 94.84% | 91.98% | 71.31% | 69.57% | 11.64% | 0.22% | 7.00% | 8.22% | 13.87% | 34.11% | |
| **SOS1-ASN-879** | 74.82% | 75.25% | 78.59% | 98.00% | 91.46% | 99.44% | 81.05% | 76.91% | 69.11% | 60.64% | 62.44% | 59.44% | 61.52% | 69.69% | 93.56% | 36.75% | 96.34% | 95.66% | 99.46% | 99.60% | 99.70% | |
| **SOS1-SER-880** | 61.74% | 68.85% | 75.13% | 75.91% | 86.83% | 87.58% | 86.24% | 87.07% | 96.52% | 98.60% | 97.26% | 97.96% | 97.30% | 87.23% | 72.15% | 4.62% | 24.05% | 36.27% | 43.16% | 50.60% | 68.07% | |
| **SOS1-SER-881** | 69.76% | 66.10% | 61.76% | 83.41% | 40.66% | 22.21% | 60.58% | 33.47% | 36.53% | 58.52% | 72.37% | 64.89% | 48.52% | 43.30% | 90.08% | 76.07% | 97.12% | 92.60% | 89.66% | 96.18% | 93.38% | |
| **SOS1-TYR-884** | 5.26% | 12.28% | 31.03% | 55.20% | 55.28% | 72.39% | 97.50% | 66.95% | 79.93% | 84.85% | 79.43% | 68.91% | 76.41% | 16.59% | 50.56% | 63.73% | 97.04% | 84.13% | 92.90% | 94.78% | 96.96% | |
| **SOS1-ARG-885** | 59.98% | 54.26% | 70.73% | 96.68% | 48.94% | 1.24% | 47.66% | 18.71% | 78.35% | 96.42% | 95.28% | 91.04% | 87.15% | 16.07% | 14.11% | 59.36% | 99.64% | 75.77% | 36.55% | 75.13% | 90.02% | |
| **SOS1-HSD-905** | 0.98% | 4.56% | 5.60% | 19.57% | 43.12% | 12.59% | 11.86% | 8.58% | 25.45% | 13.35% | 16.75% | 7.78% | 14.07% | 11.16% | 2.02% | 6.02% | 32.01% | 52.64% | 65.71% | 78.39% | 88.26% | |
| **SOS1-SER-908** | 0.40% | 4.44% | 23.41% | 38.36% | 60.54% | 93.76% | 83.63% | 99.24% | 62.32% | 43.00% | 62.63% | 44.14% | 53.26% | 68.05% | 62.40% | 9.70% | 0.58% | 2.68% | 16.83% | 29.91% | 30.73% | |
| **SOS1-GLU-909** | 0.46% | 0.18% | 10.70% | 8.86% | 27.47% | 65.31% | 36.35% | 85.09% | 45.24% | 7.96% | 13.97% | 13.27% | 22.21% | 49.30% | 43.00% | 5.74% | 2.54% | 15.65% | 8.20% | 8.60% | 2.34% | |
| **SOS1-ASP-910** | 33.95% | 18.65% | 27.29% | 27.77% | 55.64% | 44.72% | 24.55% | 66.01% | 42.42% | 36.63% | 28.27% | 17.11% | 28.29% | 61.24% | 37.17% | 9.70% | 0.28% | 13.37% | 14.83% | 17.79% | 16.27% | |
| **SOS1-HSD-911** | 95.50% | 90.54% | 96.36% | 90.54% | 97.62% | 97.92% | 85.55% | 91.06% | 96.94% | 91.44% | 96.98% | 93.38% | 93.98% | 98.24% | 98.18% | 33.97% | 2.32% | 32.33% | 99.96% | 96.26% | 99.98% | |
| **SOS1-TYR-912** | 68.47% | 62.57% | 91.24% | 59.24% | 57.46% | 91.92% | 95.80% | 96.58% | 95.08% | 95.30% | 95.30% | 95.06% | 90.18% | 93.94% | 61.06% | 11.16% | 0.00% | 14.15% | 54.36% | 32.53% | 31.49% | |
| **SOS1-LYS-913** | 20.07% | 22.77% | 36.63% | 8.38% | 8.82% | 0.70% | 1.68% | 9.14% | 9.30% | 72.15% | 87.05% | 87.47% | 82.95% | 82.03% | 86.85% | 17.73% | 0.00% | 19.03% | 87.76% | 89.44% | 95.44% | |
| **SOS1-ILE-932** | 39.76% | 14.61% | 8.64% | 0.02% | 0.02% | 12.59% | 24.37% | 66.97% | 28.85% | 7.26% | 15.11% | 2.16% | 10.76% | 16.87% | 6.24% | 3.10% | 0.00% | 9.96% | 66.09% | 89.42% | 93.36% | |
| **SOS1-LEU-934** | 1.76% | 30.53% | 69.03% | 61.46% | 55.72% | 63.95% | 45.56% | 64.09% | 24.23% | 0.04% | 0.48% | 0.32% | 0.08% | 0.08% | 0.00% | 0.00% | 0.00% | 0.00% | 0.00% | 0.00% | 0.00% | |
| **SOS1-THR-935** | 88.72% | 89.32% | 76.41% | 77.29% | 95.06% | 96.32% | 92.34% | 99.14% | 79.19% | 62.06% | 71.99% | 82.69% | 80.71% | 86.17% | 31.43% | 0.10% | 5.44% | 4.32% | 12.02% | 29.23% | 35.07% | |
| **SOS1-ASN-936** | 44.92% | 38.92% | 42.30% | 2.36% | 0.38% | 1.72% | 36.51% | 4.42% | 5.08% | 12.30% | 16.89% | 1.86% | 2.38% | 8.32% | 7.09% | 0.00% | 12.48% | 13.03% | 79.37% | 97.64% | 99.76% | |
| **SOS1-LEU-938** | 60.34% | 84.33% | 94.21% | 100.00% | 100.00% | 100.00% | 100.00% | 100.00% | 100.00% | 100.00% | 100.00% | 100.00% | 98.77% | 97.13% | 87.52% | 35.53% | 1.56% | 14.57% | 33.19% | 33.83% | 37.88% | |
| **SOS1-LYS-939** | 99.98% | 100.00% | 100.00% | 100.00% | 100.00% | 100.00% | 99.98% | 99.74% | 100.00% | 100.00% | 100.00% | 99.96% | 100.00% | 100.00% | 99.96% | 49.64% | 91.84% | 61.98% | 99.78% | 100.00% | 100.00% | |
| **SOS1-THR-940** | 2.48% | 52.22% | 75.69% | 78.17% | 76.75% | 81.81% | 77.45% | 44.96% | 97.54% | 98.04% | 98.08% | 58.44% | 32.07% | 27.95% | 57.34% | 16.85% | 0.06% | 5.30% | 9.50% | 20.83% | 51.36% | |
| **SOS1-GLU-941** | 10.40% | 20.84% | 51.48% | 99.98% | 99.98% | 99.88% | 87.17% | 63.51% | 100.00% | 100.00% | 100.00% | 100.00% | 100.00% | 100.00% | 100.00% | 67.31% | 16.09% | 73.07% | 98.18% | 93.00% | 98.78% | |
| **SOS1-GLU-942** | 69.94% | 89.18% | 98.44% | 99.20% | 99.54% | 83.09% | 75.10% | 53.25% | 61.91% | 61.41% | 65.07% | 53.10% | 50.72% | 50.50% | 44.88% | 42.30% | 17.33% | 32.33% | 72.47% | 77.97% | 78.19% | |
| **SOS1-GLY-943** | 22.51% | 31.57% | 44.56% | 43.92% | 30.49% | 8.98% | 3.38% | 0.26% | 2.56% | 7.66% | 3.36% | 0.46% | 0.52% | 0.88% | 2.28% | 71.47% | 88.48% | 91.52% | 96.38% | 96.66% | 99.44% | |
| **SOS1-ASN-944** | 33.95% | 58.80% | 67.11% | 70.37% | 33.73% | 16.61% | 1.76% | 0.74% | 2.08% | 2.06% | 0.38% | 0.12% | 0.00% | 2.14% | 7.08% | 51.36% | 79.81% | 77.33% | 84.41% | 88.84% | 86.65% | |
| **SOS1-LYS-963** | 46.72% | 68.87% | 72.45% | 20.29% | 74.01% | 83.93% | 69.05% | 56.74% | 80.31% | 40.34% | 15.65% | 9.56% | 1.16% | 6.38% | 0.72% | 0.34% | 0.00% | 7.66% | 78.83% | 51.74% | 83.85% | |
| **SOS1-GLU-966** | 9.76% | 12.69% | 3.68% | 2.66% | 22.01% | 27.59% | 36.57% | 10.44% | 24.23% | 9.34% | 0.74% | 1.80% | 0.18% | 1.68% | 0.06% | 0.00% | 0.00% | 1.80% | 10.66% | 4.52% | 25.53% | |
| **SOS1-GLU-1002** | 63.33% | 72.42% | 72.39% | 55.63% | 24.35% | 18.39% | 14.27% | 2.88% | 4.14% | 2.12% | 5.90% | 8.96% | 3.00% | 5.04% | 3.96% | 33.03% | 3.80% | 2.30% | 0.00% | 0.06% | 0.02% | |
| **SOS1-LYS-1003** | 73.27% | 81.55% | 88.58% | 86.47% | 77.99% | 77.17% | 21.01% | 25.21% | 47.66% | 63.49% | 32.55% | 57.76% | 50.16% | 71.31% | 71.59% | 57.02% | 84.01% | 60.18% | 4.64% | 8.70% | 1.20% | |
| **SOS1-THR-1006** | 29.49% | 76.03% | 80.55% | 80.41% | 38.78% | 37.23% | 29.21% | 93.74% | 95.76% | 95.08% | 86.35% | 96.08% | 95.00% | 88.80% | 68.21% | 19.13% | 0.00% | 2.50% | 1.28% | 0.48% | 0.12% | |
| | PS: Some residues with low contact frequency were not counted | | | | | | | | | | | | | | | | | | | | | |

**Figure 7.** Time-dependent contact frequency statistics: SOS1 residues contacting KRAS-G12D during the simulation (trajectory#4).

**Time-dependent contact frequency statistics: SOS1 residues contacting GDP (KRAS-G12D) during the simulation (trajectory#4)**

| | State-A | | State-B | | | | | GDP release | | | | | | | | | | | | | |
|---|---|---|---|---|---|---|---|---|---|---|---|---|---|---|---|---|---|---|---|---|---|
| **Residue** | **0-200** | **200-400** | **400-600** | **600-800** | **800-1000** | **1000-1200** | **1200-1400** | **1400-1600** | **1600-1800** | **1800-2000** | **2000-2200** | **2200-2400** | **2400-2600** | **2600-2800** | **2800-3000** | **3000-3200** | **3200-3400** | **3400-3600** | **3600-3800** | **3800-4000** | **4000-4200** |
| **SOS1-TRP-809** | 0.00% | 0.00% | 0.00% | 0.00% | 0.00% | 0.00% | 0.00% | 0.00% | 45.47% | 93.68% | 99.50% | 99.88% | 89.24% | 60.13% | 33.79% | 0.00% | 0.00% | 0.00% | 0.00% | 0.00% | 0.00% |
| **SOS1-LEU-938** | 49.30% | 88.32% | 96.77% | 99.78% | 100.00% | 100.00% | 100.00% | 100.00% | 100.00% | 100.00% | 100.00% | 100.00% | 100.00% | 95.46% | 92.94% | 85.10% | 66.63% | 55.67% | 51.44% | 45.04% | 22.13% |
| **SOS1-GLU-941** | 23.32% | 69.40% | 74.36% | 97.16% | 97.30% | 97.98% | 98.84% | 94.26% | 99.78% | 99.50% | 99.52% | 99.60% | 99.60% | 99.98% | 99.72% | 38.12% | 20.15% | 0.88% | 0.72% | 0.26% | 0.00% |
| **SOS1-GLU-942** | 45.40% | 71.37% | 83.93% | 93.55% | 95.27% | 94.29% | 98.40% | 97.98% | 99.20% | 99.26% | 87.15% | 87.25% | 84.63% | 85.29% | 79.69% | 6.74% | 0.00% | 0.00% | 0.00% | 0.00% | 0.00% |
| **SOS1-GLY-943** | 1.79% | 14.77% | 59.04% | 90.68% | 94.24% | 94.98% | 95.12% | 96.43% | 99.10% | 99.52% | 98.96% | 98.08% | 98.06% | 94.64% | 98.48% | 15.25% | 0.00% | 0.00% | 0.00% | 0.00% | 0.00% |
| **SOS1-ASN-944** | 0.00% | 0.00% | 1.84% | 10.38% | 4.92% | 1.82% | 3.08% | 14.69% | 73.13% | 86.25% | 86.71% | 85.21% | 75.45% | 75.61% | 46.84% | 3.98% | 0.00% | 0.00% | 0.00% | 0.00% | 0.00% |
| **SOS1-PRO-945** | 0.00% | 0.00% | 0.00% | 0.54% | 0.00% | 0.00% | 0.00% | 10.68% | 75.41% | 77.23% | 65.99% | 75.23% | 72.81% | 64.01% | 42.42% | 2.20% | 0.00% | 0.00% | 0.00% | 0.00% | 0.00% |
| **SOS1-GLU-946** | 0.02% | 0.00% | 0.00% | 0.04% | 0.02% | 0.12% | 0.00% | 4.72% | 41.08% | 70.85% | 81.17% | 76.73% | 63.43% | 78.31% | 56.88% | 0.06% | 0.00% | 0.00% | 0.00% | 0.00% | 0.00% |
| **SOS1-LYS-960** | 0.00% | 0.00% | 0.00% | 0.00% | 0.00% | 3.50% | 4.94% | 7.52% | 86.47% | 95.58% | 99.52% | 99.48% | 95.70% | 99.52% | 95.76% | 0.32% | 0.00% | 0.00% | 0.00% | 0.00% | 0.00% |
| | PS: Some residues with low contact frequency were not counted | | | | | | | | | | | | | | | | | | | | |

**Figure 8.** Time-dependent contact frequency statistics: SOS1 residues contacting GDP (KRAS-G12D case) during the simulation (trajectory#4).

### Time-dependent contact frequency statistics: SOS1 residues contacting KRAS-G12D during the simulation (trajectory#5)

| | State-A | State-B | | | | GDP release | | | | | | |
|---|---|---|---|---|---|---|---|---|---|---|---|---|
| **Residue** | **0-200** | **200-400** | **400-600** | **600-800** | **800-1000** | **1000-1200** | **1200-1400** | **1400-1600** | **1600-1800** | **1800-2000** | **2000-2200** | **2200-2400** |
| **PROA-GLU-589** | 0.00% | 27.65% | 21.15% | 31.75% | 44.40% | 28.06% | 23.39% | 10.79% | 3.31% | 1.85% | 0.72% | 0.78% |
| **PROA-GLU-590** | 0.54% | 15.93% | 43.46% | 5.80% | 12.77% | 30.93% | 9.24% | 8.56% | 5.74% | 7.20% | 4.76% | 1.14% |
| **PROA-ASN-591** | 0.26% | 16.26% | 17.60% | 15.82% | 27.36% | 19.23% | 24.55% | 0.18% | 0.06% | 2.74% | 0.78% | 0.28% |
| **PROA-MET-592** | 0.14% | 5.04% | 48.94% | 3.24% | 0.16% | 16.71% | 19.69% | 0.02% | 0.06% | 0.62% | 0.46% | 0.63% |
| **PROA-GLN-593** | 0.00% | 0.26% | 5.36% | 0.14% | 0.52% | 21.45% | 4.10% | 0.00% | 0.00% | 0.02% | 0.00% | 0.34% |
| **PROA-ILE-600** | 0.00% | 0.00% | 0.00% | 0.00% | 2.72% | 15.85% | 10.24% | 5.60% | 2.56% | 0.13% | 0.00% | 0.00% |
| **PROA-LYS-602** | 5.34% | 50.38% | 73.73% | 46.72% | 72.47% | 70.42% | 32.81% | 12.77% | 5.82% | 7.62% | 12.64% | 7.78% |
| **PROA-SER-807** | 73.29% | 99.54% | 90.88% | 95.75% | 90.52% | 91.52% | 40.81% | 17.94% | 1.83% | 0.43% | 0.34% | 0.37% |
| **PROA-VAL-808** | 1.32% | 6.75% | 33.65% | 33.83% | 35.32% | 35.56% | 22.36% | 0.24% | 1.62% | 0.84% | 0.71% | 1.13% |
| **PROA-TRP-809** | 73.99% | 100.00% | 99.98% | 100.00% | 100.00% | 100.00% | 70.13% | 9.98% | 7.65% | 0.78% | 0.13% | 0.83% |
| **PROA-THR-810** | 89.24% | 99.40% | 86.17% | 72.93% | 91.06% | 93.21% | 80.02% | 27.59% | 18.29% | 6.82% | 3.89% | 0.76% |
| **PROA-LYS-811** | 74.31% | 87.66% | 84.67% | 80.31% | 91.92% | 89.60% | 69.22% | 35.86% | 11.66% | 3.61% | 0.12% | 0.45% |
| **PROA-GLU-812** | 15.16% | 19.87% | 26.85% | 3.60% | 12.75% | 27.89% | 10.98% | 34.11% | 25.19% | 26.45% | 7.66% | 2.44% |
| **PROA-LYS-814** | 70.30% | 82.05% | 95.36% | 50.02% | 44.86% | 41.58% | 23.73% | 7.40% | 0.06% | 1.82% | 8.14% | 0.00% |
| **PROA-SER-876** | 75.76% | 88.89% | 91.52% | 96.96% | 81.96% | 55.90% | 49.42% | 17.84% | 7.56% | 9.34% | 3.36% | 4.04% |
| **PROA-ASN-879** | 69.44% | 78.59% | 92.32% | 99.56% | 99.28% | 89.20% | 77.14% | 67.34% | 56.42% | 59.06% | 60.66% | 57.54% |
| **PROA-SER-880** | 75.13% | 75.91% | 75.38% | 81.49% | 82.81% | 86.59% | 87.15% | 88.80% | 73.39% | 91.52% | 90.26% | 89.03% |
| **PROA-SER-881** | 70.52% | 76.02% | 62.04% | 73.35% | 86.61% | 59.32% | 67.78% | 78.66% | 63.73% | 51.60% | 70.73% | 69.62% |
| **PROA-ARG-885** | 53.68% | 50.56% | 55.54% | 52.36% | 53.10% | 64.05% | 33.25% | 53.68% | 44.36% | 45.82% | 55.02% | 46.22% |
| **PROA-SER-908** | 3.42% | 5.42% | 30.69% | 45.30% | 55.34% | 61.58% | 29.89% | 13.55% | 36.49% | 57.24% | 37.09% | 43.48% |
| **PROA-ASP-910** | 31.84% | 19.75% | 54.46% | 53.94% | 36.79% | 25.45% | 42.78% | 30.37% | 39.30% | 22.99% | 17.56% | 20.81% |
| **PROA-HSD-911** | 90.50% | 88.54% | 89.64% | 92.78% | 95.40% | 92.30% | 92.46% | 82.91% | 96.22% | 94.10% | 93.01% | 95.28% |
| **PROA-TYR-912** | 60.07% | 67.77% | 77.71% | 76.89% | 74.87% | 76.77% | 79.93% | 85.91% | 91.06% | 76.55% | 84.09% | 80.15% |
| **PROA-LYS-913** | 18.27% | 24.36% | 9.66% | 5.72% | 41.96% | 27.83% | 8.96% | 4.76% | 25.91% | 41.38% | 60.13% | 63.42% |
| **PROA-ILE-932** | 37.95% | 17.86% | 24.55% | 56.22% | 22.24% | 76.61% | 54.39% | 20.33% | 15.15% | 4.25% | 8.63% | 13.45% |
| **PROA-LEU-934** | 29.50% | 8.86% | 55.36% | 82.05% | 70.73% | 77.41% | 39.25% | 14.01% | 15.11% | 5.89% | 2.11% | 3.24% |
| **PROA-THR-935** | 85.43% | 86.02% | 97.94% | 98.82% | 99.24% | 96.46% | 93.70% | 96.72% | 97.48% | 94.40% | 85.26% | 46.78% |
| **PROA-ASN-936** | 40.73% | 37.81% | 46.32% | 43.78% | 34.21% | 37.27% | 15.25% | 10.09% | 14.81% | 9.11% | 2.01% | 3.15% |
| **PROA-LEU-938** | 88.89% | 100.00% | 100.00% | 100.00% | 100.00% | 100.00% | 100.00% | 100.00% | 100.00% | 100.00% | 100.00% | 100.00% |
| **PROA-LYS-939** | 94.13% | 100.00% | 100.00% | 100.00% | 100.00% | 99.98% | 99.80% | 99.88% | 99.92% | 99.94% | 100.00% | 98.76% |
| **PROA-THR-940** | 6.08% | 49.72% | 71.05% | 76.96% | 74.81% | 70.98% | 98.88% | 100.00% | 84.58% | 61.22% | 53.18% | 25.43% |
| **PROA-GLU-941** | 11.32% | 13.44% | 30.34% | 51.28% | 91.50% | 100.00% | 100.00% | 100.00% | 100.00% | 100.00% | 97.54% | 99.01% |
| **PROA-GLU-942** | 68.98% | 99.18% | 100.00% | 99.70% | 80.74% | 63.98% | 65.92% | 60.40% | 65.11% | 60.56% | 63.21% | 51.36% |
| **PROA-GLY-943** | 44.56% | 42.50% | 46.94% | 41.16% | 43.17% | 26.45% | 3.14% | 1.04% | 2.98% | 0.32% | 0.27% | 0.01% |
| **PROA-ASN-944** | 58.80% | 66.21% | 77.79% | 71.81% | 44.87% | 23.87% | 15.11% | 3.35% | 6.33% | 6.81% | 2.87% | 3.31% |
| **PROA-LYS-963** | 46.72% | 68.87% | 71.47% | 90.16% | 78.35% | 59.98% | 34.25% | 66.09% | 64.65% | 6.57% | 3.27% | 7.71% |
| **PROA-GLU-966** | 10.71% | 14.79% | 8.36% | 47.76% | 11.72% | 6.36% | 5.78% | 22.61% | 26.03% | 6.14% | 0.44% | 0.14% |
| **PROA-ILE-967** | 3.90% | 3.18% | 0.74% | 0.16% | 30.03% | 8.52% | 23.65% | 29.82% | 27.06% | 9.55% | 1.38% | 3.24% |
| **PROA-GLU-1002** | 73.03% | 50.12% | 38.22% | 69.47% | 71.41% | 52.77% | 40.85% | 39.62% | 12.34% | 4.63% | 2.78% | 5.43% |
| **PROA-LYS-1003** | 70.01% | 78.85% | 60.08% | 64.71% | 80.91% | 80.63% | 85.27% | 86.55% | 58.62% | 72.61% | 44.63% | 36.59% |
| **PROA-THR-1006** | 66.03% | 79.49% | 88.05% | 64.79% | 77.81% | 85.13% | 77.61% | 76.59% | 86.08% | 85.05% | 76.24% | 70.21% |
| | PS: Some residues with low contact frequency were not counted | | | | | | | | | | | |

**Figure 9.** Time-dependent contact frequency statistics: SOS1 residues contacting KRAS-G12D during the simulation (trajectory#5).

**Time-dependent contact frequency statistics: SOS1 residues contacting GDP (KRAS-G12D) during the simulation (trajectory#5)**

| | State-A | State-B | | | | GDP release | | | | | | | |
|---|---|---|---|---|---|---|---|---|---|---|---|---|---|
| Residue | 0-200 | 200-400 | 400-600 | 600-800 | 800-1000 | 1000-1200 | 1200-1400 | 1400-1600 | 1600-1800 | 1800-2000 | 2000-2200 | 2200-2400 | ... |
| SOS1-TRP-809 | 0.00% | 0.00% | 0.00% | 0.00% | 0.00% | 0.00% | 25.47% | 30.47% | 45.47% | 93.68% | 97.50% | 88.88% | |
| SOS1-LEU-938 | 80.12% | 100.00% | 100.00% | 100.00% | 100.00% | 100.00% | 100.00% | 100.00% | 100.00% | 100.00% | 99.94% | 99.56% | |
| SOS1-GLU-941 | 69.40% | 97.98% | 98.74% | 89.84% | 87.72% | 96.08% | 99.76% | 99.70% | 99.70% | 99.34% | 99.54% | 99.62% | |
| SOS1-GLU-942 | 79.26% | 99.20% | 99.40% | 99.22% | 99.40% | 67.23% | 93.80% | 85.41% | 64.55% | 84.37% | 74.55% | 83.46% | |
| SOS1-GLY-943 | 2.36% | 15.15% | 13.96% | 21.04% | 90.32% | 97.24% | 99.52% | 97.74% | 91.18% | 94.48% | 94.12% | 95.65% | |
| SOS1-ASN-944 | 0.00% | 0.00% | 0.00% | 0.00% | 0.00% | 2.58% | 33.87% | 32.85% | 52.19% | 65.08% | 74.18% | 88.10% | |
| SOS1-PRO-945 | 0.00% | 0.00% | 0.00% | 0.00% | 0.00% | 1.44% | 34.79% | 21.37% | 46.77% | 53.54% | 70.12% | 67.86% | |
| SOS1-GLU-946 | 0.02% | 0.00% | 0.00% | 0.00% | 0.02% | 0.34% | 26.65% | 27.19% | 48.78% | 63.04% | 67.34% | 71.20% | |
| SOS1-LYS-960 | 0.00% | 0.00% | 0.00% | 0.00% | 0.00% | 0.32% | 27.73% | 21.69% | 61.32% | 83.44% | 88.12% | 86.54% | |
| | PS: Some residues with low contact frequency were not counted | | | | | | | | | | | | |

**Figure 10.** Time-dependent contact frequency statistics: SOS1 residues contacting GDP (KRAS-G12D case) during the simulation (trajectory#5).

### Time-dependent contact frequency statistics: SOS1 residues contacting KRAS-G12D during the simulation (trajectory#6)

| | State-A | | State-B | | GDP release | | | | | | | | |
|---|---|---|---|---|---|---|---|---|---|---|---|---|---|
| **Residue** | **0-200** | **200-400** | **400-600** | **600-800** | **800-1000** | **1000-1200** | **1200-1400** | **1400-1600** | **1600-1800** | **1800-2000** | **2000-2200** | **2200-2400** | **......** |
| **PROA-GLU-589** | 0% | 23.78% | 40.22% | 18.81% | 25.65% | 20.12% | 9.83% | 5.44% | 1.76% | 1.43% | 0.43% | 0.71% | |
| **PROA-GLU-590** | 4.14% | 10.78% | 12.75% | 0% | 0.52% | 3.92% | 2.18% | 5.31% | 0.13% | 0.00% | 3.61% | 0.20% | |
| **PROA-ASN-591** | 1.67% | 7.06% | 8.38% | 0.18% | 2.58% | 4.06% | 1.02% | 0.49% | 0.00% | 0.68% | 2.12% | 0.44% | |
| **PROA-MET-592** | 1.38% | 6.42% | 21.75% | 0.68% | 2.18% | 11.27% | 7.02% | 10.19% | 0.84% | 13.45% | 10.29% | 5.60% | |
| **PROA-GLN-593** | 0% | 0.26% | 2.06% | 2.18% | 17.68% | 38.13% | 43.64% | 37.11% | 1.36% | 9.54% | 17.19% | 7.89% | |
| **PROA-LYS-595** | 0% | 0% | 0.06% | 0.04% | 12.22% | 22.69% | 11.78% | 3.46% | 4.56% | 1.71% | 3.58% | 0.12% | |
| **PROA-LYS-602** | 1.24% | 52.98% | 71.39% | 40.10% | 43.76% | 40.12% | 35.73% | 15.68% | 3.41% | 10.73% | 15.71% | 10.18% | |
| **PROA-SER-807** | 75.41% | 97.31% | 83.34% | 84.6% | 87.54% | 45.13% | 18.72% | 2.91% | 0.03% | 0.1% | 0.31% | 0.37% | |
| **PROA-VAL-808** | 0.12% | 5.70% | 32.26% | 41.04% | 41.98% | 25.71% | 20.43% | 3.72% | 0.05% | 1.18% | 0.51% | 0.46% | |
| **PROA-TRP-809** | 74.51% | 100.00% | 99.90% | 99.98% | 98.56% | 63.01% | 16.06% | 8.31% | 0.72% | 2.94% | 0.00% | 0.00% | |
| **PROA-THR-810** | 85.35% | 98.01% | 90.66% | 98.17% | 98.75% | 84.68% | 50.02% | 10.71% | 4.32% | 0.78% | 1.02% | 0.13% | |
| **PROA-LYS-811** | 75.68% | 85.46% | 74.4% | 82.28% | 60.56% | 31.07% | 20.12% | 9.82% | 2.38% | 1.68% | 0.02% | 0.33% | |
| **PROA-GLU-812** | 17.01% | 18.24% | 41.61% | 52.49% | 19.66% | 16.02% | 13.44% | 0.78% | 0.23% | 1.58% | 0.38% | 0.53% | |
| **PROA-LYS-814** | 85.36% | 86.35% | 90.48% | 78.03% | 63.25% | 48.02% | 36.16% | 12.02% | 1.04% | 2.84% | 0.74% | 0.00% | |
| **PROA-SER-876** | 76.14% | 78.74% | 79.27% | 99.60% | 89.74% | 53.40% | 47.70% | 35.16% | 14.84% | 1.98% | 1.31% | 4.57% | |
| **PROA-ASN-879** | 78.32% | 80.08% | 93.78% | 99.50% | 89.92% | 73.15% | 73.01% | 67.02% | 59.82% | 61.46% | 54.08% | 62.02% | |
| **PROA-SER-880** | 70.56% | 67.02% | 67.80% | 93.80% | 90.90% | 85.34% | 86.12% | 90.14% | 86.14% | 86.42% | 77.34% | 80.42% | |
| **PROA-SER-881** | 79.04% | 56.00% | 39.02% | 60.20% | 43.38% | 35.40% | 56.42% | 70.08% | 60.64% | 50.72% | 63.31% | 76.58% | |
| **PROA-TYR-884** | 13.06% | 31.03% | 54.79% | 71.18% | 78.28% | 79.01% | 82.92% | 77.24% | 68.03% | 77.12% | 66.38% | 51.76% | |
| **PROA-ARG-885** | 60.78% | 57.86% | 62.59% | 41.86% | 60.20% | 46.56% | 37.75% | 78.35% | 86.47% | 84.36% | 36.48% | 20.17% | |
| **PROA-HSD-905** | 6.57% | 27.96% | 36.05% | 22.8% | 14.03% | 26.34% | 17.64% | 16.61% | 10.58% | 15.27% | 11.66% | 7.32% | |
| **PROA-SER-908** | 4.54% | 8.42% | 27.17% | 49.47% | 83.24% | 60.36% | 45.32% | 60.01% | 42.84% | 54.06% | 67.37% | 57.43% | |
| **PROA-ASP-910** | 36.87% | 23.04% | 35.79% | 59.58% | 67.37% | 43.38% | 37.59% | 30.44% | 19.61% | 26.45% | 22.54% | 38.04% | |
| **PROA-HSD-911** | 94.52% | 89.76% | 96.50% | 96.54% | 98.00% | 95.83% | 90.55% | 95.74% | 92.62% | 90.18% | 97.68% | 94.46% | |
| **PROA-TYR-912** | 69.34% | 94.06% | 61.51% | 88.34% | 92.74% | 94.16% | 93.48% | 94.32% | 94.15% | 93.46% | 92.76% | 65.41% | |
| **PROA-LYS-913** | 25.34% | 27.87% | 35.53% | 9.64% | 4.25% | 9.68% | 70.35% | 85.22% | 86.37% | 80.74% | 83.01% | 84.65% | |
| **PROA-LEU-934** | 3.43% | 31.06% | 71.25% | 79.56% | 74.56% | 34.62% | 10.82% | 3.31% | 1.35% | 0.34% | 0.81% | 0.00% | |
| **PROA-THR-935** | 90.02% | 95.46% | 99.22% | 99.38% | 99.98% | 78.12% | 64.26% | 73.09% | 81.79% | 79.76% | 83.28% | 35.46% | |
| **PROA-ASN-936** | 46.02% | 40.22% | 10.36% | 48.02% | 9.92% | 7.28% | 9.32% | 13.76% | 3.02% | 4.06% | 7.34% | 8.47% | |
| **PROA-LEU-938** | 80.98% | 99.02% | 100.00% | 100.00% | 100.00% | 100.00% | 100.00% | 100.00% | 98.76% | 96.64% | 97.83% | 85.36% | |
| **PROA-LYS-939** | 98.78% | 100.00% | 100.00% | 99.82% | 99.44% | 100.00% | 100.00% | 100.00% | 97.06% | 98.76% | 100.00% | 96.79% | |
| **PROA-THR-940** | 6.88% | 55.82% | 48.76% | 52.42% | 74.2% | 74.68% | 88.25% | 78.88% | 55.64% | 34.50% | 21.03% | 10.41% | |
| **PROA-GLU-941** | 13.04% | 41.05% | 94.78% | 93.39% | 88.98% | 71.72% | 100.00% | 100.00% | 98.76% | 100.00% | 99.46% | 100.00% | |
| **PROA-GLU-942** | 79.94% | 99.18% | 99.76% | 88.74% | 78.31% | 60.21% | 63.08% | 59.27% | 50.10% | 55.68% | 53.58% | 42.64% | |
| **PROA-GLY-943** | 46.86% | 22.51% | 46.97% | 45.16% | 13.69% | 10.86% | 7.04% | 4.14% | 0.34% | 1.46% | 0.98% | 1.14% | |
| **PROA-ASN-944** | 55.83% | 67.11% | 73.84% | 78.19% | 42.44% | 24.13% | 6.71% | 4.86% | 2.34% | 2.66% | 0.43% | 1.03% | |
| **PROA-LYS-963** | 47.94% | 70.87% | 82.79% | 82.71% | 88.48% | 65.35% | 58.24% | 78.41% | 32.33% | 9.67% | 3.76% | 2.36% | |
| **PROA-GLU-966** | 10.04% | 13.58% | 28.39% | 16.51% | 32.97% | 22.17% | 15.06% | 0.74% | 1.80% | 0.18% | 1.68% | 0.06% | |
| **PROA-GLU-1002** | 83.33% | 79.04% | 79.79% | 62.66% | 40.58% | 25.46% | 14.47% | 10.87% | 4.78% | 4.32% | 1.32% | 2.39% | |
| **PROA-LYS-1003** | 76.43% | 84.76% | 87.17% | 88.85% | 59.47% | 54.73% | 61.34% | 34.65% | 57.80% | 45.36% | 69.33% | 72.78% | |
| **PROA-THR-1006** | 46.23% | 79.99% | 84.32% | 80.85% | 79.79% | 85.36% | 94.28% | 84.75% | 95.35% | 90.76% | 86.38% | 70.01% | |
| | PS: Some residues with low contact frequency were not counted | | | | | | | | | | | | |

**Figure 11.** Time-dependent contact frequency statistics: SOS1 residues contacting KRAS-G12D during the simulation (trajectory#6).

**Time-dependent contact frequency statistics: SOS1 residues contacting GDP (KRAS-G12D) during the simulation (trajectory#6)**

| | State-A | | State-B | | GDP release | | | | | | | |
|---|---|---|---|---|---|---|---|---|---|---|---|---|
| **Residue** | **0-200** | **200-400** | **400-600** | **600-800** | **800-1000** | **1000-1200** | **1200-1400** | **1400-1600** | **1600-1800** | **1800-2000** | **2000-2200** | **2200-2400** |
| **SOS1-TRP-809** | 0.00% | 0.00% | 0.00% | 0.00% | 0.00% | 0.00% | 40.07% | 90.58% | 97.51% | 98.98% | 80.43% | 75.21% |
| **SOS1-LEU-938** | 85.41% | 100.00% | 100.00% | 100.00% | 100.00% | 99.89% | 98.79% | 99.34% | 100.00% | 98.53% | 96.73% | 95.43% |
| **SOS1-GLU-941** | 79.40% | 90.44% | 97.90% | 95.90% | 95.78% | 94.08% | 95.51% | 96.62% | 95.36% | 97.61% | 96.48% | 89.12% |
| **SOS1-GLU-942** | 69.78% | 90.32% | 99.66% | 85.93% | 44.24% | 89.02% | 94.37% | 85.35% | 83.23% | 84.93% | 82.39% | 78.34% |
| **SOS1-GLY-943** | 6.78% | 17.85% | 14.40% | 55.16% | 94.68% | 90.13% | 93.57% | 91.78% | 95.38% | 93.76% | 89.69% | 85.78% |
| **SOS1-ASN-944** | 0.00% | 0.00% | 0.00% | 0.14% | 7.36% | 14.69% | 67.93% | 80.73% | 83.29% | 75.39% | 75.78% | 74.74% |
| **SOS1-PRO-945** | 0.00% | 0.00% | 0.00% | 0.02% | 0.32% | 10.68% | 70.31% | 78.03% | 75.49% | 72.71% | 69.68% | 68.76% |
| **SOS1-GLU-946** | 0.02% | 0.00% | 0.00% | 0.00% | 0.00% | 5.02% | 39.88% | 69.75% | 78.54% | 65.93% | 79.01% | 63.58% |
| **SOS1-LYS-960** | 0.00% | 0.00% | 0.00% | 0.12% | 0.84% | 17.64% | 75.97% | 90.68% | 95.42% | 94.98% | 96.73% | 89.98% |
| | PS: Some residues with low contact frequency were not counted | | | | | | | | | | | |

**Figure 12.** Time-dependent contact frequency statistics: SOS1 residues contacting GDP (KRAS-G12D case) during the simulation (trajectory#6).
